# Time-Resolved Interactome Profiling Deconvolutes Secretory Protein Quality Control Dynamics

**DOI:** 10.1101/2022.09.04.506558

**Authors:** Madison T. Wright, Bibek Timalsina, Valeria Garcia Lopez, Jake Hermanson, Sarah Garcia, Lars Plate

## Abstract

Many cellular processes are governed by protein-protein interactions that require tight spatial and temporal regulation. Accordingly, it is necessary to understand the dynamics of these interactions to fully comprehend and elucidate cellular processes and pathological disease states. To map de novo protein-protein interactions with time-resolution at an organelle-wide scale we developed a quantitative mass-spectrometry method, time-resolved interactome profiling (TRIP). We apply TRIP to elucidate aberrant protein interaction dynamics that lead to the protein misfolding disease congenital hypothyroidism. We deconvolute altered temporal interactions of the thyroid hormone precursor thyroglobulin with pathways implicated in hypothyroidism pathophysiology such as Hsp70/90 assisted folding, disulfide/redox processing, and N-glycosylation. Functional siRNA screening identified VCP and TEX264 as key protein degradation components whose inhibition selectively rescues mutant prohormone secretion. Ultimately, our results provide novel insight into the temporal coordination of protein homeostasis, and our TRIP method should find broad applications in investigating protein folding diseases and cellular processes.

## INTRODUCTION

Protein-protein interactions drive functional diversity within cells and are often closely connected to the observed phenotypes for cellular processes and disease states (Bludau & Aebersold, 2020). Protein homeostasis (proteostasis) is a critical cellular process that relies on tightly regulated protein interactions. The proteostasis network (PN), consisting of protein folding chaperones, trafficking, and degradation components, maintains the integrity of the proteome by ensuring the appropriate trafficking and localization of properly folded proteins while recognizing misfolded, potentially detrimental states and routing them for degradation (Karagöz *et al*, 2019; Needham *et al*, 2019; Behnke *et al*, 2016; Pohl & Dikic, 2019). The concerted action of hundreds of proteostasis factors is referred to as protein quality control (PQC). Perturbations to the PN through genetic, age-related, or environmental factors manifest in several disease states, including amyloidosis, neurodegeneration, cancer, and others (Taldone *et al*, 2020; McDonald *et al*, 2022; Wright *et al*, 2021; Kuo *et al*, 2021; Marinko *et al*, 2021).

Identifying and quantifying protein-protein interactions has been critical for comprehending the pathogenesis of these disease states. Approaches include yeast two-hybrid systems, co-immunoprecipitation coupled with western blot analysis, the Luminescence-based Mammalian IntERactome (LUMIER) assay, as well as affinity purification – mass spectrometry (AP-MS) (Taipale *et al*, 2012, 2014; Piette *et al*, 2021; Rizzolo *et al*, 2017; Rizzolo & Houry, 2019; Wright & Plate, 2021). These methods have been powerful for mapping steady-state proteostasis interactions to disease states, yet most lack the ability to measure interaction dynamics over time. Proximity labeling mass spectrometry (BioID & APEX-MS) has had limited use to spatiotemporally resolve protein-protein interactions only following protein maturation, as synchronization of newly synthesized protein populations is challenging to achieve (Lobingier *et al*, 2017; Perez Verdaguer *et al*, 2022). Unnatural amino acid incorporation methods, such as biorthogonal non-canonical amino acid tagging (BONCAT), pulsed azidohomoalanine (PALM), or heavy isotope labeled azidohomoalanine (HILAQ) can identify newly synthesized proteins but have not focused on a single endogenously expressed protein or group of proteins in the context of disease (Dieterich *et al*, 2006; Bagert *et al*, 2014; Ma *et al*, 2018; McClatchy *et al*, 2015; Ma *et al*, 2017; Howden *et al*, 2013; van Bergen *et al*, 2022).

Similar DNA and RNA labeling approaches are used to temporally resolve nascent DNA-protein and RNA-protein interactions in cell culture and whole organisms. Isolation of proteins on nascent DNA (iPOND) has revealed the timing of DNA-protein interactions during replication and chromatin assembly (Cortez, 2017; Sirbu *et al*, 2011; Munden *et al*, 2022). Thiouracil crosslinking mass spectrometry (TUX-MS) and viral cross-linking and solid-phase purification (VIR-CLASP) are used to study the timing of RNA-protein interactions during viral infection (Kim *et al*, 2020; Phillips *et al*, 2016). In contrast, no methods were previously available to identify de novo proteinprotein interactions of newly synthesized proteins in a client specific manner with temporal resolution, which has motivated our efforts here.

In earlier work, we mapped the interactome of the secreted thyroid prohormone thyroglobulin (Tg) comparing the WT protein to secretion-defective mutations implicated in congenital hypothyroidism (CH) (Wright *et al*, 2021). Tg is a heavily post-translationally modified, 330 kDa prohormone that is necessary to produce triiodothyronine (T3) and thyroxine (T4) thyroid specific hormones (Citterio *et al*, 2019; Coscia *et al*, 2020). Tg biogenesis relies extensively on distinct interactions with the PN to facilitate folding and eventual secretion. Our previous results identified topological changes in Tg-PN interactions among CH-associated Tg mutants compared to WT. Nonetheless, the lack of temporal resolution precludes more mechanistic discernment of these changes in PN interactions. While some changes may simply correlate with disease pathogenesis, others may be directly responsible for the aberrant PQC and secretion defect of the mutant Tg variants.

To address this shortcoming, we developed time-resolved interactome profiling (TRIP) to capture and quantify interactions between Tg and interacting partners throughout the life cycle of the protein. We found that Tg mutants are characterized by both discrete changes with select PN components and broad temporal alterations across Hsp70/90, N-glycosylation, and disulfide/redox processing pathways. Moreover, we find that these perturbations are correlated with alterations in interactions with degradation components. We coupled our TRIP method with functional siRNA screening and uncovered that VCP (p97) and TEX264 are two key regulators of Tg processing. VCP and TEX264 inhibition or silencing in thyroid cells rescued the secretion of mutant Tg, representing – to our knowledge – the first restorative approach based upon proteostasis modulation to increase mutant Tg secretion.

## RESULTS

### TRIP temporally resolves Tg interactions with PQC components

To develop the time-resolved interactome profiling method, we envisioned a two-stage enrichment strategy utilizing epitope-tagged immunoprecipitation coupled with pulsed biorthogonal unnatural amino acid labeling and functionalization (**Fig 1A**). Cells are pulse-labeled with homopropargylglycine (Hpg) to synchronize newly synthesized protein populations. After the Hpg pulse, samples are collected across time points throughout a chase period (**Fig 1A**, Box 1) (Kiick *et al*, 2001; Beatty *et al*, 2006). The Hpg alkyne incorporated into the newly synthesized population of protein is then conjugated to biotin using copper-catalyzed alkyne-azide cycloaddition (CuAAC) (**Fig 1A**, Box 2). Subsequently, the first stage of the enrichment strategy globally captures the client protein and binding partners using epitope-tagged immunoprecipitation, followed by elution (**Fig 1A**, Box 3). The second enrichment step then utilizes a biotin-streptavidin pulldown to capture the Hpg pulse-labeled, and CuAAC conjugated population, enriching samples into time-resolved fractions (**Fig 1A**, Box 4) (Li *et al*, 2020; Thompson *et al*, 2019).

**Figure 1.**
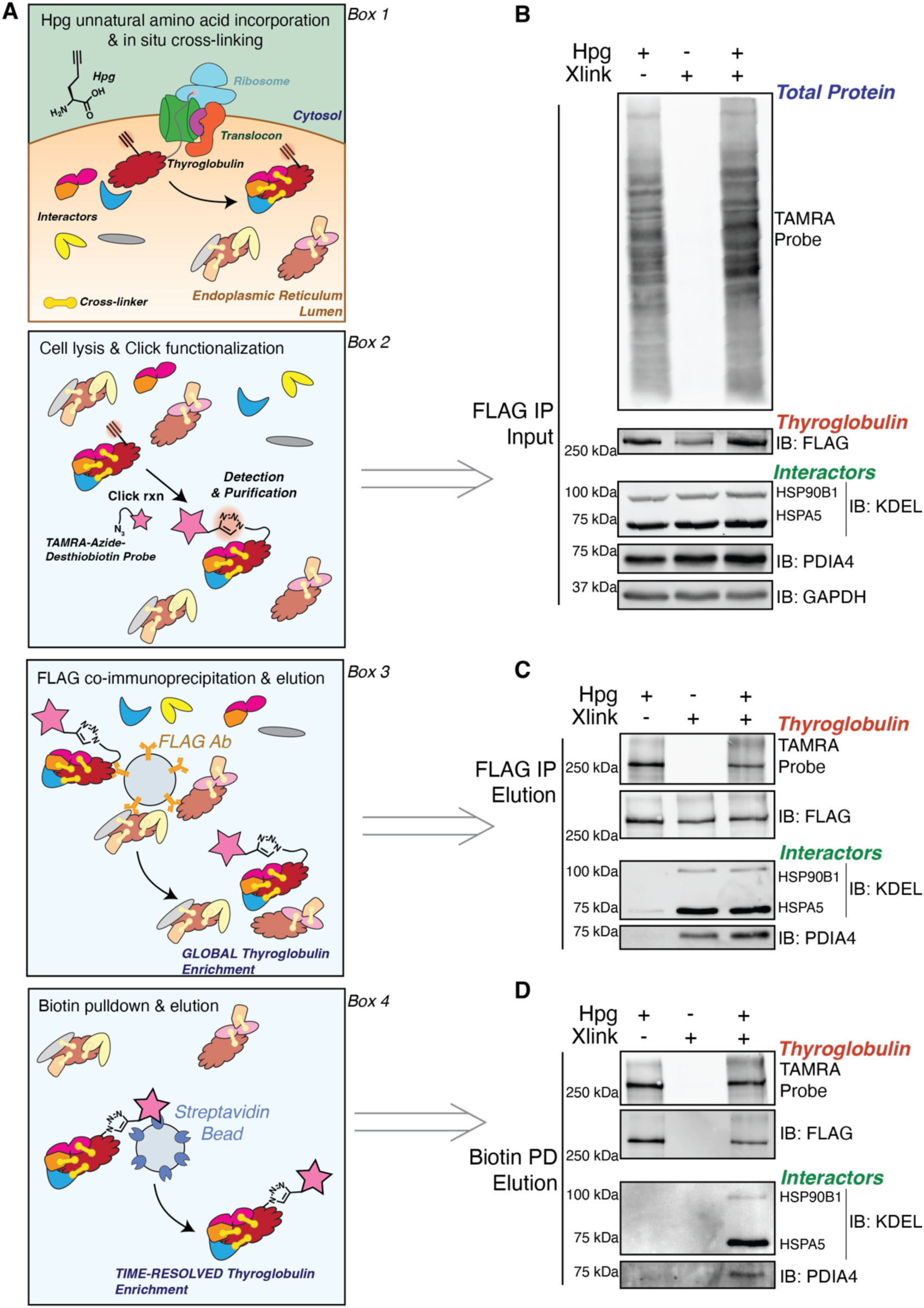
Time-Resolved Interactome Profiling to Identify Interactions with newly synthesized Proteins. (A) Key steps necessary for the TRIP workflow. *Box 1* shows pulsed unnatural amino acid labeling with Hpg to incorporate and alkyne functional group into newly synthesized proteins followed by in-situ cross-linking of protein interactions using DSP. *Box 2* shows labeled protein functionalization with copper catalyzed azide-alkyne cycloaddition (CuAAC) click chemistry. *Box 3* shows FLAG co-immunoprecipitation to globally enrich labeled and unlabeled Tg. *Box 4* shows subsequent streptavidin-biotin enrichment of the pulse-labeled, time-resolved Tg fractions. (B-D) Western blot analysis of the two-stage purification strategy with continuous Hpg labeling. FRT cells stably expressing WT Tg were continuously labeled with Hpg (200μM) for 4 h and crosslinked with DSP (0.5mM) for 10 minutes to capture transient proteostasis network interactions (+ Xlink). Lysates were functionalized with a TAMRA-Azide-PEG-Desthiobiotin probe using copper catalyzed azide-alkyne cycloaddition (CuAAC) and subjected to the dual affinity purification scheme as described. (B) FLAG IP inputs as described in *Box 2* in (A). Top shows fluorescence image of TAMRA labeled proteins, and bottom shows immunoblots of Tg (IB: FLAG) and interactors HSP90B1, HSPA5, PDIA4 and loading control GAPDH. (C) FLAG IP elutions as described in *Box 3* in (A). Top shows fluorescence image of TAMRA labeled Tg and total Tg (IB: FLAG). Bottom shows immunoblots of interactors (HSP90B1, HSPA5, and PDIA4). Samples were then subjected to biotin pulldown. (D) Biotin pulldown elutions as described in Box 4 in (A). Top shows fluorescence image of rhodamine labeled Tg and total Tg (IB: FLAG). Bottom shows immunoblots of interactors (HSP90B1, HSPA5, and PDIA4).

Thyroglobulin was chosen as the model secretory client protein. We generated isogenic Fischer rat thyroid (FRT) cells that stably expressed FLAG-tagged Tg (Tg-FT), including WT or mutant variants (A2234D and C1264R) (**Fig EV1**). WT Tg is readily secreted from the FRT cells, while C1264R Tg shows only minimal residual secretion (∼2% of WT when detected after immunoprecipitation) (**Fig EV1D**). A2234D is fully secretion deficient, consistent with prior characterization of these variants (Hishinuma *et al*, 1999; Pardo *et al*, 2009). We first set out to determine if Tg could tolerate immunoprecipitation after pulse Hpg labeling, dithiobis(succinimidyl propionate) (DSP) crosslinking, and CuAAC conjugation with a trifunctional tetramethylrhodamine (TAMRA)-Azide-Polyethylene Glycol (PEG)-Desthiobiotin probe. We showed previously that a C-terminal FLAG-tag is tolerated by Tg and allows efficient immunoprecipitation, while DSP crosslinker aids in capturing transient protein-protein interactions that take place during Tg processing (Wright *et al*, 2021). We pulse-labeled FRT cells expressing WT or C1264R Tg for 4 h with Hpg, performed DSP crosslinking, CuAAC, and tested our two-stage enrichment strategy via western blot analysis (**Fig 1B–D**). Pulsed Hpg labeling, DSP crosslinking, and CuAAC did not significantly affect immunoprecipitation efficiency and allowed for robust two-stage enrichment of WT and C1264R Tg-FT with well validated Tg interactors HSPA5 (BiP), HSP90B1 (Grp94), and PDIA4 (ERp72) (**Fig 1B,C**) (Menon *et al*, 2007; Baryshev *et al*; Wright *et al*, 2021). Furthermore, the C-terminal FLAG-tag and Hpg labeling are necessary for this two-stage enrichment strategy, and DSP crosslinking is necessary to capture these interactions after stringent wash steps (**Fig 1D**, **Fig EV2**).

Next, we investigated whether TRIP could temporally resolve interactions with these PN components. We pulse labeled WT Tg FRT cells with Hpg for 1 h, followed by a 3 h chase in regular media capturing time points in 30-minute intervals and analyzing via western blot or TMTpro LC-MS/MS (**Fig 2A**). Our previous study indicated that ∼70% of WT Tg-FT was secreted after 4 hours, while approximately 50% of A2234D and 15% of C1264R was degraded after the same time period (Wright *et al*, 2021). Therefore, we reasoned that a 3 h chase period would be enough time to capture the majority of Tg interactions throughout processing, secretion, cellular retention, and degradation, while still being able to capture an appreciable amount of sample for analysis. For WT Tg, interactions with HSPA5 peaked within the first 30 minutes of the chase period and rapidly declined, in line with previous observations, but PDIA4 interactions were not detectable by western blot analysis (**Fig 2B**) (Menon *et al*, 2007; Kim & Arvan, 1995). To test the ability of TRIP to distinguish temporal differences in PQC interactions across mutant Tg variants, we performed TRIP on FRT cells expressing C1264R Tg, a known patient mutation implicated in CH (Hishinuma *et al*, 1999; Kanou *et al*, 2007). For C1264R, interactions with HSPA5 were highly abundant at the 0 h time point and remained mostly steady throughout the first 1.5 h (**Fig 2C**). A similar temporal profile was also observed for HSP90B1. Additionally, interactions with PDIA4 were detectable for C1264R and were found to gradually increase throughout the first 1.5 h of the chase period, before rapidly declining (**Fig 2C**). We noticed similar temporal profiles for PDIA4 and HSPA5 to our western blot analysis, when measured via TMTpro LC-MS/MS as further outlined below (**Fig 2D-E**). In particular, the HSPA5 WT Tg interaction declined within the first hours, yet for C1264R Tg, the HSPA5 interactions remained mostly steady over the 3-hour chase period. (**Fig 2E**).

**Figure 2.**
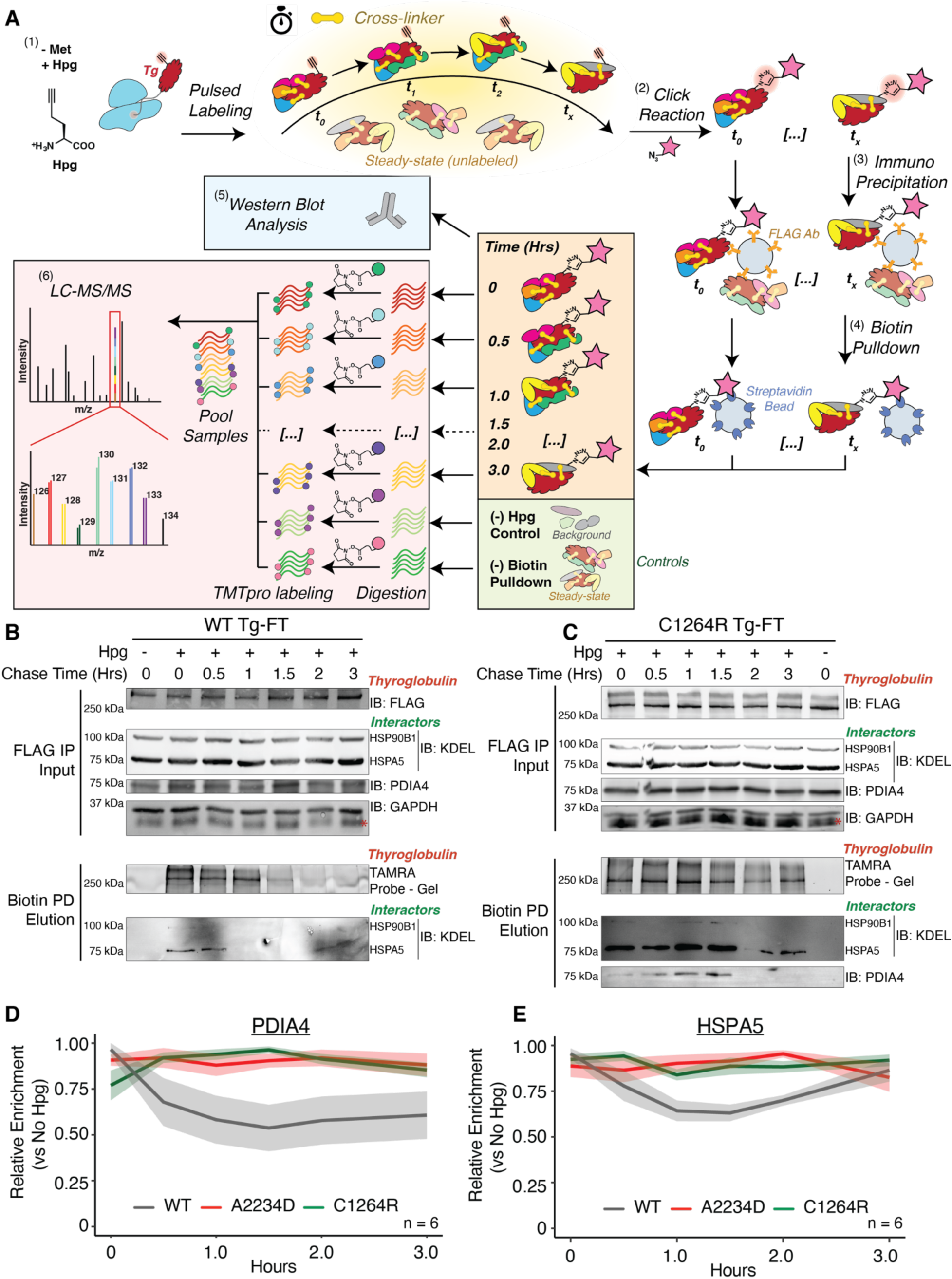
TRIP Temporally Resolves Tg Interactions with Protein Quality Control Components. (A) Workflow for TRIP protocol utilizing western blot or mass spectrometric analysis of time-resolved interactomes. (1) Cells are pulse-labeled with Hpg (200μM final concentration) for 1 h, chased in regular media for specified time points, and cross-linked with DSP (0.5mM) for 10 minutes to capture transient proteoastasis network interactions; (2) Lysates are functionalized with a TAMRA-Azide-PEG-Desthiobiotin probe using copper CuAAC Click reaction; (3) Lysates undergo the first stage of the enrichment strategy where the Tg-FT is globally captured and enriched using immunoprecipitation; (4) Eluted Tg-FT populations from the global immunoprecipitation undergo biotin-streptavidin pulldown to capture the pulse Hpg-labeled, and CuAAC conjugated population of Tg-FT, enriching samples into time-resolved fractions; (5) Time-resolved fraction may then undergo western blot analysis or (6) quantitative liquid chromatography – tandem mass spectrometry (LC-MS/MS) analysis with tandem mass tag (TMTpro) multiplexing or analysis. The (-) Hpg control channel is used to identify enriched interactors and a (-) Biotin pulldown channel to act as a booster (or carrier). (B and C) TRIP western blot analysis of WT Tg-FT (B) and C1264R Tg-FT (C). Samples were processed as described above in (A). FLAG IP Input panel shows immunoblot of Tg (IB: 1FLAG), and interactors HSP90B1, HSPA5, PDIA4 and loading control GAPDH. Biotin pulldown elution panel shows fluorescence image of TAMRA labeled Tg and immunoblot of Tg interactors HSP90B1 and HSPA5. Validated Tg interactors show higher and delayed enrichment with the misfolded C1264R Tg mutant in (C) compared to WT in (B). PDIA4 interactions were not detectable by western blot analysis for biotin pulldown elution with WT Tg. (D and E) Plots showing the scaled enrichment of select Tg interactors HSPA5 and PDIA4 compared across constructs. Samples were processed as described above in (A) and analyzed by mass spectrometry. Solid line corresponds to mean and shading represents the SEM (N = 5 for WT Tg; N = 6 for A2234D and C1264R Tg). Data in **Dataset EV 4**.

These data highlight the utility of TRIP to not only identify changes in protein interactions over time but also monitor how these interactions differ for a given protein of interest across disease states. Moreover, these data corroborate our previous findings that steady-state interactions with HSPA5, HSP90B1 and PDIA4 are elevated for C1264R Tg (Wright *et al*, 2021).

### TRIP identifies altered temporal dynamics associated with Tg processing

We benchmarked the utility of our TRIP approach to temporally resolve previously identified and novel interactors, as the Tg interactome has not been fully characterized in native tissue. We focused on A2234D and C1264R Tg as they present with distinct defects in Tg processing and mutations are localized in separate structural domains (Kanou *et al*, 2007; Hishinuma *et al*, 1999). Following the Hpg pulse-chase labeling scheme and dual affinity purification, the time-resolved Tg fractions were trypsin/Lys-C digested and labeled individually with isobaric TMTpro tags. Subsequently, two sets of TRIP time course samples (0, 0.5, 1, 1.5, 2, and 3 h) could be pooled using the 16plex TMTpro and analyzed by LC-MS/MS (**Fig 2A**). In total, 5 biological replicates were analyzed for WT and 6 biological replicates were analyzed for A2234D and C1264R (**Table EV3**). Aside from the experimental samples, we utilized a (–) biotin pulldown booster (or carrier) channel with cells that were pulse-labeled with Hpg, underwent CuAAC functionalization, and immunoprecipitation, but did not undergo biotin-streptavidin enrichment (**Fig 2A**). This booster sample acted to 1) aid in increased peptide/protein identification – compared to the much lower abundant chase samples; and 2) benchmark Tg interactors in FRT cells compared to our previously published dataset (Tsai *et al*, 2020; Petelski *et al*, 2021). When comparing the booster channel to our (–)Hpg negative control samples, most previously identified Tg interactors were strongly and significantly enriched (**Fig EV3**). Our dataset in this study showed appreciable overlap between our previous results in HEK293T cells identifying 75 of the previous 171 Tg interactors and identifying 198 new interactors (**Fig 3A**). Several ribosomal and proteasomal subunits, trafficking factors, and lysosomal components were not identified in our previous dataset (**Fig EV3, Dataset EV2**). We then took our list of previously identified interactors and PQC components found to be enriched in the (–) biotin pulldown samples compared to (–) Hpg and carried these proteins forward to time-resolved analysis utilizing the Hpg-chase samples (**Table EV1**).

**Figure 3.**
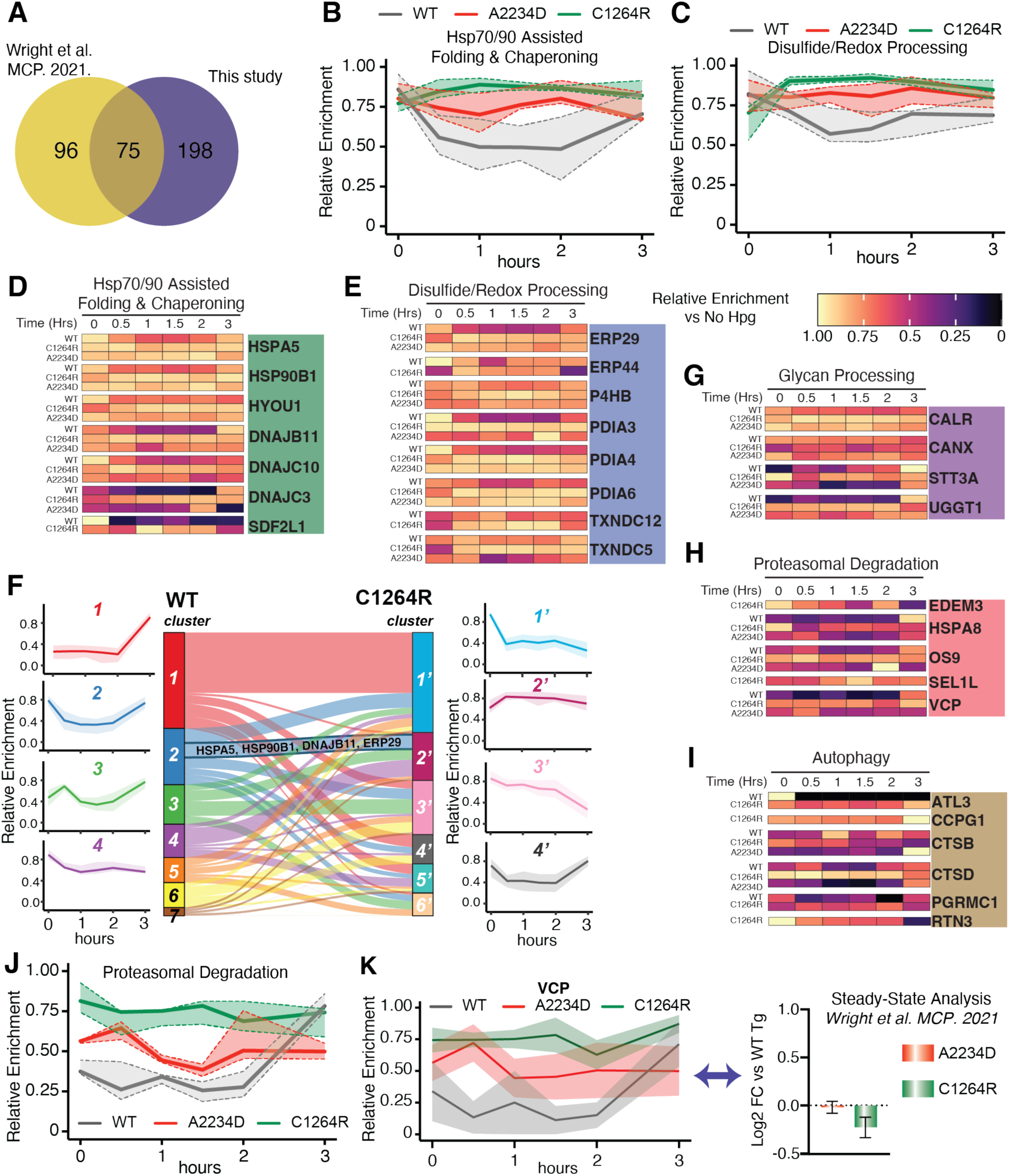
TRIP Identifies Altered Temporal Dynamics Associated with Tg Processing. (A) Venn diagram showing the overlap in proteostasis components identified as Tg interactors here compared to our previous data set (Wright *et al*, 2021). (-) biotin pulldown vs (-) Hpg samples were used to identify Tg interactors in FRT cells. Data available in **Dataset EV3**. (B-C) Plot showing the relative pathway enrichment for Hsp70/90 assisted folding & chaperoning interactors (B) and disulfide/redox processing interactors (C) with WT, A2234D, or C1264R Tg constructs. For this and all subsequent panels, the average log2 fold change enrichment value across time points for a given interactor were used to scale data. Positive enrichment was scaled from 0 to 1. Lines represent the median scaled enrichment for the group of interactors and shades correspond to the first and third quartile range. All source data for this and subsequent panels can be found in **Dataset EV4**. (D-E) Heatmap showing the relative enrichment for individual Hsp70/90 assisted folding & chaperoning interactors (D) and disulfide/redox processing interactors (E) with WT, A2234D, or C1264R Tg constructs. Relative enrichment scaled as described above in (C). (F) Unbiased k-means clustering of TRIP profiles to determine co-regulated groups of interactors. Aggregate time profiles for the most prominent clusters are shown on the left (WT) and the right (C1264R). The line corresponds to the mean scaled log2 fold enrichment and the shading represents the 25 – 75 % quartile range within each cluster. The Sankey plot in the center shows the shift of interactors between clusters from WT to C1264R. (G-I) Heatmap showing the relative enrichment for select, for select glycan processing (G), proteasomal degradation (H) and autophagy interactors (I) with WT, A2234D, or C1264R Tg. (J) Plot showing the relative pathway enrichment for proteasomal degradation interactors. (K) Plot comparing the relative enrichment of VCP throughout the time course for WT, C1264R, and A2234D Tg. The TRIP data (left) is contrasted to aggregate (steady-state) interacomics data (right) (Wright *et al*, 2021). TRIP resolves dynamic VCP interaction changes with mutant Tg, while these changes are muted in the aggregate data.

To map the time-resolved Tg PN interactome changes, we considered the Hpg chase samples (0 – 3 h) and compared the enrichment of Tg interactors to the (–) Hpg control. The enrichments were normalized to Tg protein levels to account for gradual changes due to Tg secretion or degradation, and positive enrichment values were scaled from 0 to 1. We organized the interactors according to distinct PN pathways known to influence Tg processing (**Appendix Fig S1, Datasets EV3,EV4**).

To benchmark the TRIP methodology, we chose to monitor a set of well-validated Tg interactors and compare the time-resolved PN interactome changes to our previously published steady-state interactomics dataset (Wright *et al*, 2021). Previously, we found that CALR (Calreticulin), CANX (Calnexin), ERP29 (PDIA9), ERP44, and P4HB interactions with mutants A2234D or C1264R Tg exhibited little to no change when compared to WT under steady-state conditions (**Fig EV4A**). However, in our TRIP dataset we were able to uncover distinct temporal changes in engagement that were previously masked within the steady-state data. Our time-resolved data deconvolutes these aggregate measurements, revealing prolonged CALR, ERP29, and P4HB engagements for both A2234D and C1264R Tg mutants compared to WT (**Fig EV4B-F**). We found that these measurements for key interactors and PN pathways exhibited robust reproducibility, as exemplified by the standard error of the mean for the TRIP data (**Fig EV4B-I, Appendix Fig S1B**).

Next, we monitored temporal changes more broadly across proteostasis network pathways. We found that both A2234D and C1264R exhibited prolonged interactions with components of Hsp70/90 and disulfide/redox processing pathways (**Fig 3B-E**). Particularly, A2234D and C1264R showed increased interactions with HSPA5, HSP90B1, HYOU1 (Grp170), DNAJC3 (ERdj6), and SDF2L1 throughout the chase period (**Fig 3D**). Conversely, WT interactions peaked at the 0 h time point and consistently tapered off for many of these components (**Fig 3C**). Similarly divergent temporal interactions were observed in the case of disulfide/redox processing components. PDIA3 (ERp57), PDIA4, PDIA6 (ERp5), and ERP29 have all been heavily implicated in Tg processing (**Fig 3E**) (di Jeso *et al*, 2005; Menon *et al*, 2007; di Jeso *et al*, 2014; Baryshev *et al*, 2006). While WT interactions with these components showed similar trends as Hsp70/90 chaperoning components – peaking at the 0 h time point and consistently decreasing – prolonged mutant interactions peaked later at 1-1.5 h for both A2234D and C1264R (**Fig 3C,E**). Furthermore, individual protein disulfide isomerase interactions peaked at distinct times, thereby revealing an order to their engagement (**Fig EV4G-I**).

To assess temporal interaction changes in an unbiased fashion and identify protein groups exhibiting comparative behavior, we carried out k-means clustering of the temporal profiles for WT and C1264R. This analysis revealed a large divergence in the interaction profiles. For WT Tg, only one cluster exhibited steadily decreasing interactions (cluster 4), while others increased with time, or showed peaks at intermediate time points (**Fig 3F, Fig EV5A**). On the other hand, C1264R largely exhibited clusters with decreasing interactions over time (**Fig 3F, Fig EV5B**). Cluster 2 for WT with bimodal interactions at early and late time points contains many Hsp70/90 chaperoning components. For C1264R Tg, many Hsp70/90 chaperoning components and disulfide/redox-processing components are instead part of cluster 2’, which exhibited an initial rise in interactions strength before plateauing (**Fig 3F, Fig EV5A,B**). This divergent temporal engagement between WT Tg and the destabilized C1264R mutant is aligned with the patterns observed in the manual grouping (**Fig 3B,C**), highlighting that the unbiased temporal clustering can reveal broader patterns in the reorganization of the proteostasis dynamics.

### TRIP highlights link between glycan processing and ER-phagy pathways

One area of particular interest was the crosstalk and correlation between interactions with glycan processing components and degradation pathways. The link between the glycosylation state of ER clients and ER-associated degradation (ERAD) is well established, whereas more recently defined autophagy at the ER (ER-phagy) or ER to lysosome-associated degradation (ERLAD) represent alternative degradation mechanisms for ER clients (Christianson *et al*, 2008, 2012; Fregno *et al*, 2021; Chiritoiu *et al*, 2020; Fregno *et al*, 2018). Previously, we showed that A2234D and C1264R differ in interactions with N glycosylation components, particularly the oligosaccharyltransferase (OST) complex. Efficient A2234D degradation required both STT3A and STT3B isoforms of the OST, which mediate co-translational or post-translational N-glycosylation, respectively (Kelleher *et al*, 2003; Cherepanova & Gilmore, 2016). TRIP revealed differential interactions with glycosylation components that may lead to altered degradation dynamics. Many glycan processing enzymes, lectin chaperones, and several subunits of the OST complex were identified (**Fig 3G**, **Appendix Fig S1A**). While STT3A interactions across all constructs showed similar temporal profiles, we observed prolonged interactions for lectin chaperones CALR and CANX with mutant Tg (**Fig EV4B,C**). The most striking difference observed was with the “gatekeeping” glycosyltransferase UGGT1 (**Fig 3G**). This protein regulates glycoprotein folding through the CANX/CALR cycle by re-glycosylating ER clients, and thus triggering reengagement with the lectin chaperone cycle (Lamriben *et al*, 2016). UGGT1 interactions with WT remain moderate from 0.5-1.5 hand peak at 3 h, while interactions for A2234D and C1264R peaked earlier and were more pronounced throughout the chase period. These differential interactions with UGGT1 may suggest changes in monitoring of the Tg folded state. Moreover, CANX has been directly linked to emerging mechanisms of ERLAD for glycosylated ER clients (Forrester *et al*, 2019; Fregno *et al*, 2018).

Proteasomal and autophagic degradation pathways exhibited broad differences in interaction dynamics for WT and mutant Tg (**Fig 3H-J**). The ERAD-associated lectin OS9 (Erlec2), and ATPase VCP both peaked at the 3 h chase time point for WT Tg (**Fig 3H,K**). Conversely, A2234D exhibited more prolonged VCP interactions, and OS9 interactions peaked at the 2 h time point (**Fig 3H,K**). For C1264R, we observed much stronger and prolonged VCP interactions, as well as additional interactions with the ERAD-associated mannosidase EDEM3 and E3 ubiquitin ligase adaptor SEL1L (Hrd3) (Christianson *et al*, 2008, 2012) (**Fig 3H**). Most notably, our previous aggregate steady-state data showed no significant difference for VCP interactions between WT and mutant Tg, yet our TRIP workflow was able to resolve these temporal dynamics (**Fig 3K**) (Wright *et al*, 2021). Further highlighting the utility of this novel TRIP methodology.

Our original dataset identified several lysosomal components, yet it was unclear how Tg might be delivered to the lysosome. Recent work has identified several selective ER-phagy receptors, and highlighted ER-phagy mechanisms for the clearance of mutant prohormones and other destabilized clients from the ER (Chen *et al*, 2021; Cunningham *et al*, 2019). It was intriguing then to identify several lysosomal and autophagy-related components and observe differential temporal profiles across WT and C1264R Tg constructs (**Fig 3I**, **Appendix Fig S1**). For A2234D, interactions with these components were sparser. The most intriguing observation was the enrichment of three different ER-phagy receptors, ATL3 (Atlastin-3), CCPG1 (Cpr8), and RTN3 (Reticulon 3) between WT and C1264R, along with the RTN3 adaptor protein PGRMC1 (**Fig 3I**) (Chen *et al*, 2019; Liang *et al*, 2018; Smith *et al*, 2018; Grumati *et al*, 2017; Chen *et al*, 2021). CCPG1 and RTN3 were found to specifically interact with C1264R, with RTN3 interactions peaking at 0 h and then decreasing, while CCPG1 interactions peaked later (**Fig 3I**). In the C1264R k-means clustered profiles, autophagy interactions largely group together in the same cluster, showing stronger interactions at earlier time points. In the same cluster are glycosylation components (UGGT1,STT3B, and MLEC), further supporting a possible coordination for C1264R Tg between lectin-dependent protein quality control and targeting to autophagy (**Fig EV5B,C**).

### siRNA screening discovers key regulators of construct specific Tg processing

We developed an RNA interference screening platform to investigate whether the temporal interaction changes discovered by TRIP are functionally important for Tg PQC. Moreover, we were interested in identifying factors whose modulation may act to rescue mutant Tg secretion. HEK293 cells were engineered to stably express nanoluciferase-tagged Tg constructs (Tg-NLuc) and screened against 167 Tg interactors and related PN components (see **Dataset EV5** for the list of genes). The NLuc tag allowed us to monitor changes to both intracellular Tg abundance in the cell lysates, and Tg levels in the conditioned media to assess secretion rates in a 96-well format (**Fig 4A**). Importantly, the NLuc-tag did not alter the secretion of WT Tg, and CH-associated mutants maintained the same secretion deficiency (**Fig EV6**) (England *et al*, 2016). Silencing of NAPA (α-SNAP) and LMAN1 (Ergic53) were found to increase WT Tg-NLuc lysate abundance but had no effects on the two mutants (**Fig 4B, Appendix Fig S2A, Dataset EV5**). NAPA is a member of the Soluble N-ethylmaleimide-sensitive factor Attachment Protein (SNAP) family and plays a critical role in vesicle fusion and docking, while LMAN1 is a mannose-specific lectin that functions as a glycoprotein cargo receptor for ER-to-Golgi trafficking (Song *et al*, 2017; Zhao *et al*, 2007; Marinko *et al*, 2021). For mutant Tg-NLuc constructs we found CTSZ (cathepsin Z) silencing decreased A2234D Tg-NLuc lysate abundance, while GUSB (β-glucoronidase) silencing increased C1264R lysate abundance (**Fig 4B**).

**Figure 4.**
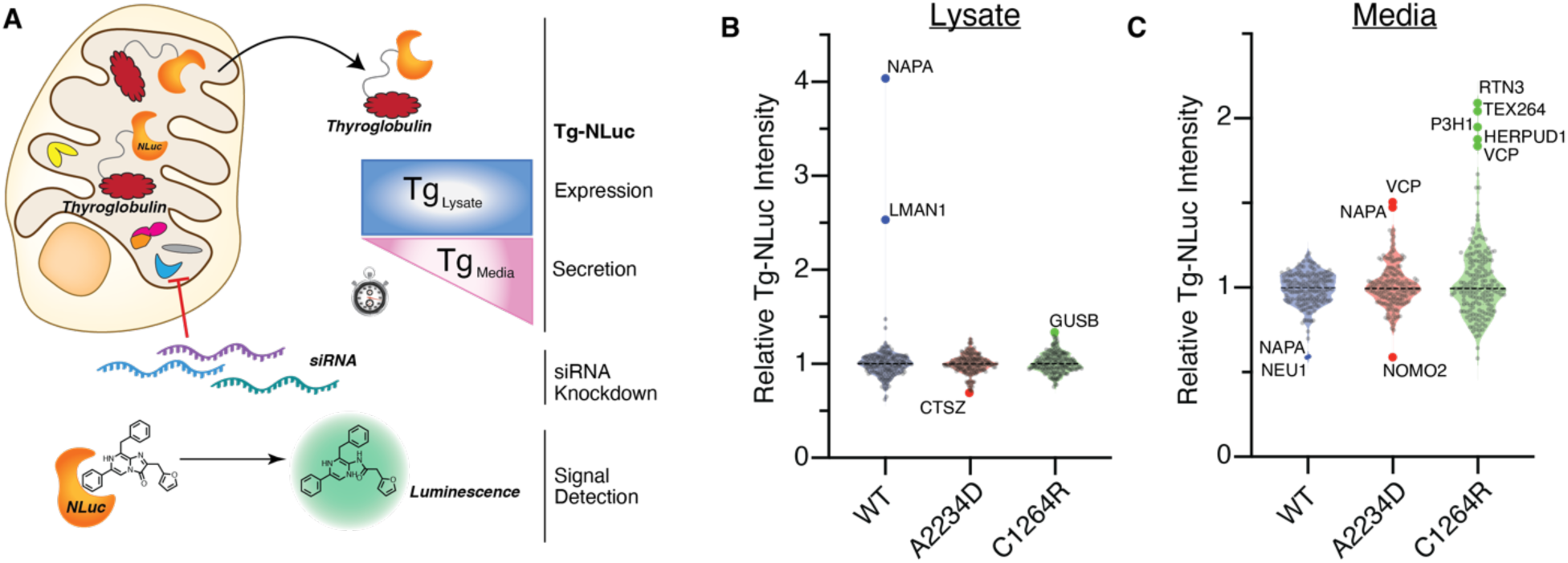
siRNA Screening Finds Regulators of Construct Specific Tg Processing. (A) siRNA screening workflow utilizing NLuc-tagged Tg to monitor lysate and media abundance. Approximately 36 hafter transfection with 25nM siRNAs cells were replenished with fresh media and Tg-NLuc abundance in lysate and media was measured after 4 husing the nano-glo luciferase assay system. (B) Violin plots showing the relative Tg-NLuc abundance changes in lysate with siRNA knockdown of select genes. Tg-NLuc abundance in lysate was measured 4 h after replenishing with fresh media using the nano-glo luciferase assay system. Hits labeled within the plot were defined as changes in Tg-NLuc abundance by greater than 3σ. Full data available in **Appendix Fig S2** and **Dataset EV5**. N = 2 for WT and A2234D Tg. N = 3 for C1264R Tg. (C) Violin plots showing the relative Tg-NLuc abundance changes in media with siRNA knockdown of select genes. Sample processing and cutoff criteria to identify hits are described above in (B). N = 2 for WT and A2234D Tg. N = 3 for C1264R Tg.

Remarkably, we identified 6 genes whose silencing rescued mutant Tg-NLuc secretion in a construct specific manner. NAPA silencing increased secretion of A2234D Tg-NLuc (**Fig 4C**). This contrast to the reduction in WT secretion with NAPA silencing may suggest an alternative role for NAPA in regulating mutant Tg processing as other proteins involved in vesicular fusion and trafficking have been implicated in ER-phagy (Cui *et al*, 2019; Liang *et al*, 2020). Silencing of P3H1 (Lepre1), an ER-resident prolyl hydroxylase, increased C1264R Tg-NLuc secretion but not WT nor A2234D (**Fig 4C**) (Vranka *et al*, 2004). Silencing of several protein degradation genes robustly increased mutant Tg secretion: VCP, HERPUD1, TEX264 and RTN3. VCP silencing increased both A2234D and C1264R Tg-NLuc secretion (**Fig 4C**). VCP is associated with ERAD but also aids in several diverse cellular functions including the interplay between proteasomal and autophagic degradation (Hill *et al*, 2021; Christianson *et al*, 2008, 2012). VCP silencing exclusively affecting mutant Tg corroborates our TRIP dataset, and suggest a more prominent role for VCP in mutant Tg PQC compared to WT. VCP interactions were sparse for WT Tg while they remained more steady throughout the chase period for the mutants (**Fig 3H,K**). HERPUD1, TEX264 and RTN3 silencing selectively increased C1264R secretion, but did not alter WT nor A2234D secretion (**Fig 4C**). HERPUD1 is a ubiquitin-like protein and associates with VCP during ERAD (Christianson *et al*, 2012; Needham *et al*, 2019; Okuda-Shimizu & Hendershot, 2007). The ER-phagy receptors RTN3 and TEX264 localize to sub domains of the ER to facilitate degradation of specific ER clients and organellular regions (Chino *et al*, 2019; Fielden *et al*, 2022; An *et al*, 2019; Grumati *et al*, 2017; Chen *et al*, 2021; Cunningham *et al*, 2019). Unfortunately, TEX264 was not identified in our TRIP data, but RTN3 was found to specifically interact with only C1264R (**Fig 3I**, **Appendix Fig S2**). Of note, while the ER-phagy receptor CCPG1 was identified in our mass spectrometry dataset siRNA silencing of CCPG1 did not significantly alter Tg-NLuc abundance in lysate or media, nor did silencing of SEC62 or RETREG1 (FAM134B), two additional ER-phagy receptors found to regulate ER dynamics (**Appendix Fig S2**) (Fumagalli *et al*, 2016; Bhaskara *et al*, 2019).

This is the first study to broadly investigate the functional implications of Tg interactors and other PQC network components on Tg processing. Coupling these data with our TRIP methodology helped to deconvolute PQC dynamics associated with Tg and identify pathways implicated in the aberrant secretion of CH-associated Tg mutations. The discovery of several protein degradation components as hits for rescuing mutant Tg secretion may suggest that the blockage of degradation pathways can broadly rescue the secretion of A2234D and C1264R mutant Tg, a phenomenon similarly found for destabilized CFTR implicated in the protein folding disease cystic fibrosis (Vij *et al*, 2006; Pankow *et al*, 2015; McDonald *et al*, 2022).

### Trafficking and degradation factors selectively regulate Tg processing in thyroid cells

We examined the hits from the initial siRNA screen in FRT cells stably expressing Tg constructs to test whether their silencing exhibited similar phenotypes in thyroid specific tissue. Thyroid tissue must synthesize and fold a large amount of Tg as it is the main protein produced and can make up more than 50% of all protein components within the thyroid gland (di Jeso & Arvan, 2016). Silencing of NAPA led to a ∼50% increase in WT Tg lysate abundance, while NAPA and LMAN1 silencing both led to marginal decreases in WT Tg secretion after 4 h, consistent with the results in HEK293 cells (**Fig 5A,B**; **Fig EV7A,B**). Using ^35^S pulse-chase analysis, we confirmed that NAPA silencing significantly increased lysate retention by 15% over 4 h and decreased secretion by 18% (**Fig EV7H**). To understand if these results were directly attributable to NAPA function, we performed complementation experiments where FRT cells treated with NAPA siRNAs were co-transfected with a human NAPA plasmid. WT Tg lysate abundance decreased when NAPA expression was complemented, confirming that the observed retention phenotype could be attributed to NAPA silencing (**Fig EV7I**). These results established that NAPA acts as a prosecretion factor for WT Tg.

**Figure 5.**
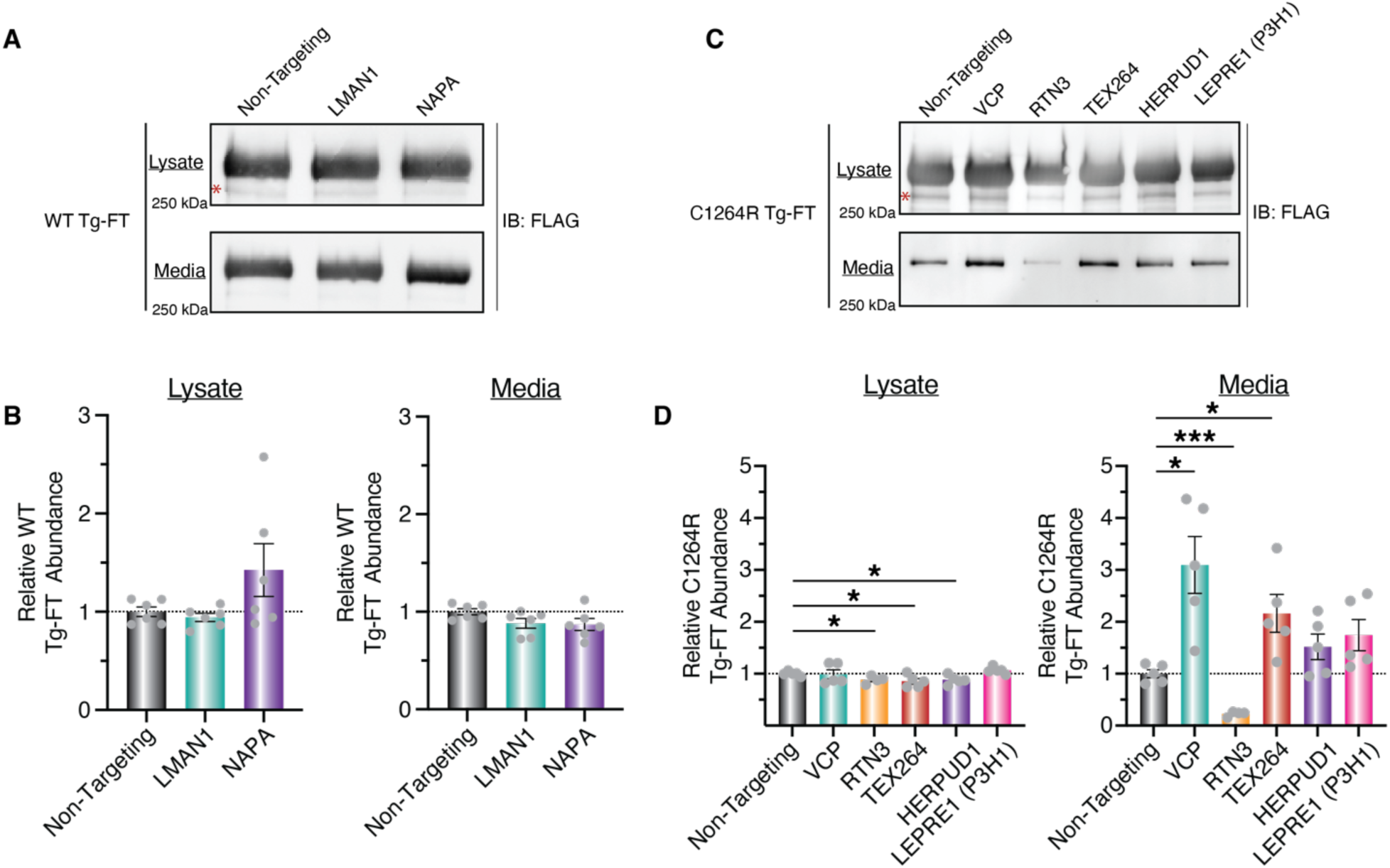
Trafficking and Degradation Factors Regulate Tg Processing in FRT cells. (A and B) Western blot analysis (A) and quantification (B) of WT Tg-FT secretion from FRT cells transfected with select siRNAs hits from initial screening data set. Red asterisk denotes a nonspecific background band within the western blot. Cells were transfected with 25nM siRNAs for 36 h, media exchanged and conditions for 4 h, Tg-FT was immunoprecipitated from lysate and media samples, and Tg-FT amounts were analyzed via immunoblotting. N = 6. (C and D) Western blot analysis (C) and quantification (D) of C1264R Tg-FT secretion from FRT cells transfected with select siRNA hits from the initial screening data set. Red asterisk denotes a non-specific background band within the western blot. Cells were transfected with 25nM siR-NAs for 36 h, media exchanged and conditions for 8 h, Tg-FT was immunoprecipitated from lysate and media samples, and Tg-FT amounts were analyzed via immunoblotting. All statistical testing performed using an unpaired student’s t-test with Welch’s correction. *p<0.05, **p<0.005, ***p<0.0005. N = 5 (one RTN3 sample excluded due to sample handling error).

In C1264R Tg FRT cells, RTN3, TEX264, and HERPUD1 silencing marginally, yet significantly decreased C1264R lysate abundance (**Fig 5C,D**, **Fig EV7C-G**). Moreover, VCP and TEX264 silencing significantly increased C1264R secretion by 3-fold and 2-fold respectively after 8 h, in line with our results in HEK293 cells (**Fig 5C,D**). In contrast, silencing of RTN3 significantly decreased C1264R Tg secretion by 5-fold, in opposition to the increased secretion observed in HEK293 cells. Several individual VCP and TEX264 siRNAs were able to partially recapitulate these increased secretion phenotype on C1264R Tg-FT, confirming that the effect is mediated by the respective gene silencing (**Fig EV7J,K**).

### Mutant Tg is selectively enriched with the ER-phagy receptor TEX264

Intrigued by the finding that TEX264 silencing increased C1264R Tg secretion without affecting WT Tg, we asked whether TEX264 exhibited differential interactions with Tg constructs. HEK293 Tg-NLuc cells were transfected with either a fluorescent GFP control, or C-terminal FLAG-tagged ER-phagy receptors, followed by FLAG co-IP. The NLuc assay and Western blotting was then used to monitor Tg enrichment (**Fig 6A**). The negative control SEC62 did not yield any appreciable enrichment of Tg compared to GFP control (**Fig 6B**), consistent with SEC62 not impacting Tg constructs in the siRNA screen. Conversely, co-IP of TEX264 resulted in the enrichment of all Tg variants compared to GFP control when monitored by NLuc assay, with C1264R and A2234D being significantly enriched. C1264R Tg exhibited a 3-fold increased interaction compared to WT Tg and almost a 2-fold increase compared to A2234D Tg (**Fig 6B**). This increase in C1264R enrichment was also observable by western blot analysis (**Fig 6C**). Additionally, we monitored Tg enrichment with ER-phagy receptors CCPG1 and RTN3 via Western blot as both were found to be C1264R Tg interactors within our TRIP dataset. RTN3L is found to be the only RTN3 isoform involved in ER turnover via ER-phagy (Grumati et al, 2017). WT and C1264R Tg-NLuc were modestly enriched with RTN3L compared to control samples expressing GFP. Conversely, we found that all Tg variants exhibited modest interactions with CCPG1 compared to control samples expressing GFP, although less than with TEX264 (**Fig EV8**).

**Figure 6.**
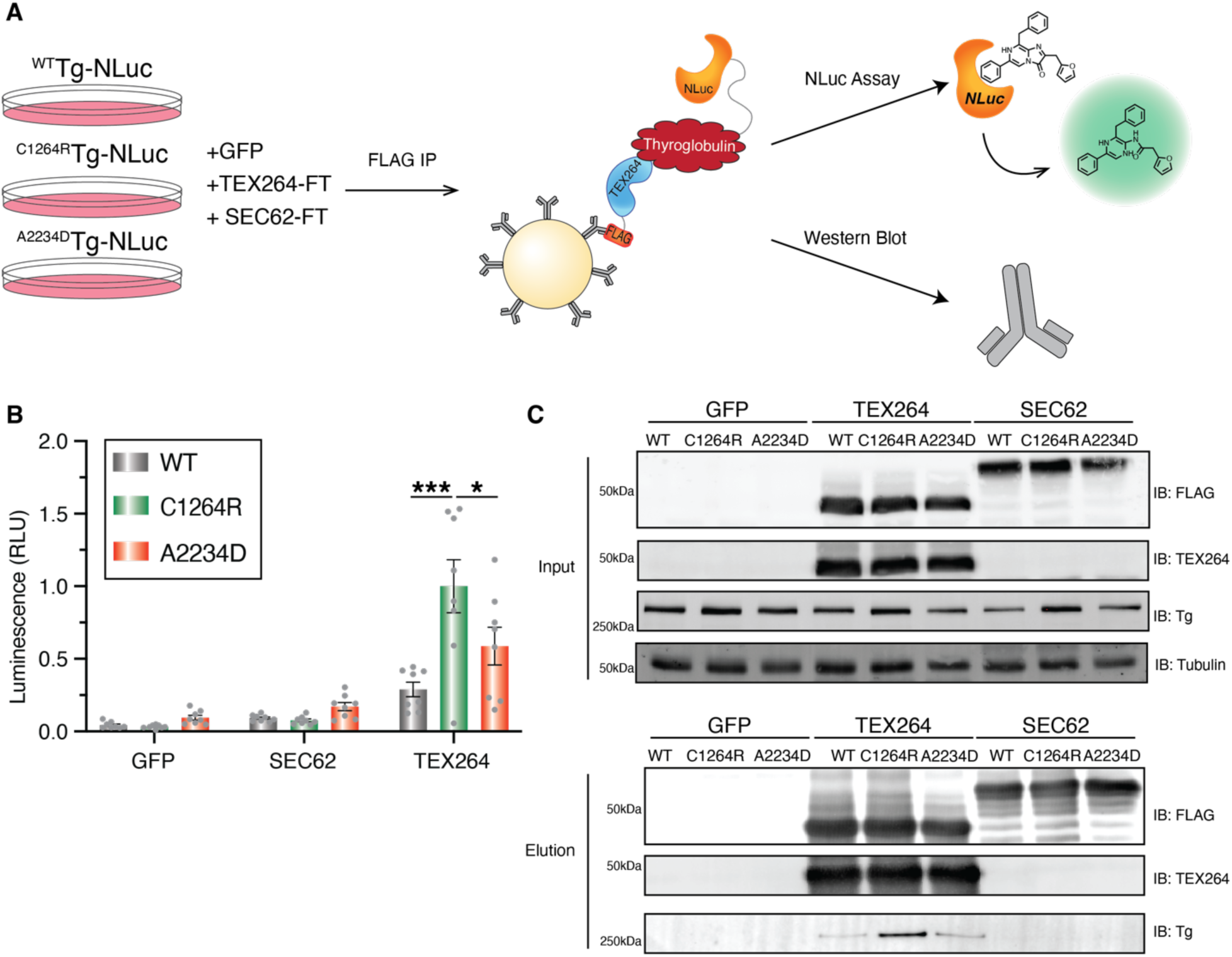
Mutant Tg is Selectively Enriched with the ER-phagy Receptor TEX264. (A) Schematic for western blot and NLuc analysis for identifying TEX264-Tg interactions. HEK293T cells stably expression WT or mutant Tg-NLuc were transfected with FLAG-tagged ER-phagy receptors TEX264, SEC62, or GFP control. After FLAG Co-IP, samples were analyzed using the using the nano-glo luciferase assay system or immunoblot analysis to monitor Tg enrichment. (B) Luminescence data from FLAG-CoIP and nano-glo luciferase analysis. Tg is selectively enriched with TEX264 compared to GFP fluorescent control, or SEC62. Mutant Tg exhibits higher enrichment compared to WT. Data represented as mean ± SEM. Statistical testing was performed using a one-way ANOVA with post hoc Tukey’s multiple testing corrections, *p<0.05, **p<0.005, ***p<0.0005. N = 8. (C) Western blot analysis of FLAG Co-IP samples. Top panel shows IP inputs and bottom panel IP elutions with Tg (IB:Tg), ER-phagy receptors (IB:FLAG and IB:TEX264) and loading control Tubulin. Tg is selectively enriched with TEX264 compared to GFP fluorescent control, or SEC62, with C1264R Tg exhibiting higher enrichment compared to WT.

Together, these data suggest that TEX264, CCPG1, or RTN3L engage with Tg during processing, and CH-associated Tg mutants may be selectively targeted to TEX264. Furthermore, ER-phagy may be considered as a degradative pathway in Tg processing, as other studies have mainly focused on Tg degradation through ERAD (Tokunaga *et al*, 2000; Menon *et al*, 2007).

### Pharmacological VCP Inhibition Selectively Rescues C1264R Tg Secretion

Considering the promising finding that silencing of degradation factors by siRNA rescued C1264R secretion, we sought to investigate whether pharmacological modulation of select Tg processing components could similarly improve Tg secretion. There are no selective inhibitors currently available for TEX264, but several for VCP. We monitored C1264R Tg secretion from FRT cells in the presence of VCP inhibitors ML-240, CB-5083, and NMS-873. ML-240 and CB-5083 are ATP-competitive inhibitors that preferentially target the D2 domain of VCP subunits, whereas NMS-873 is a non-ATP-competitive allosteric inhibitor which binds at the D1-D2 interface of VCP subunits (Chou *et al*, 2013, 2014; Anderson *et al*, 2015; le Moigne *et al*, 2017; Tang *et al*, 2019). ML-240 and NMS-873 have been shown to decrease both proteasomal degradation and autophagy, in line with VCP playing a role in both processes (Chou *et al*, 2013, 2014; Her *et al*, 2016). Conversely, while CB-5083 is known to decrease proteasomal degradation it has been shown to increase autophagy. (Anderson *et al*, 2015; le Moigne *et al*, 2017; Tang *et al*, 2019). We found that treatment with ML-240 was able to significantly increase C1264R Tg secretion without altering C1264R Tg lysate abundance, corroborating the siRNA silencing data (**Fig 7A,B**). To investigate whether this rescue of C1264R Tg secretion with ML-240 treatment was specific to C1264R Tg, we also monitored WT Tg abundance. In contrast, we observed that ML-240 significantly reduced WT Tg abundance in both lysate and media 4 and 8 hafter treatments (**Fig 7C,D**).

**Figure 7.**
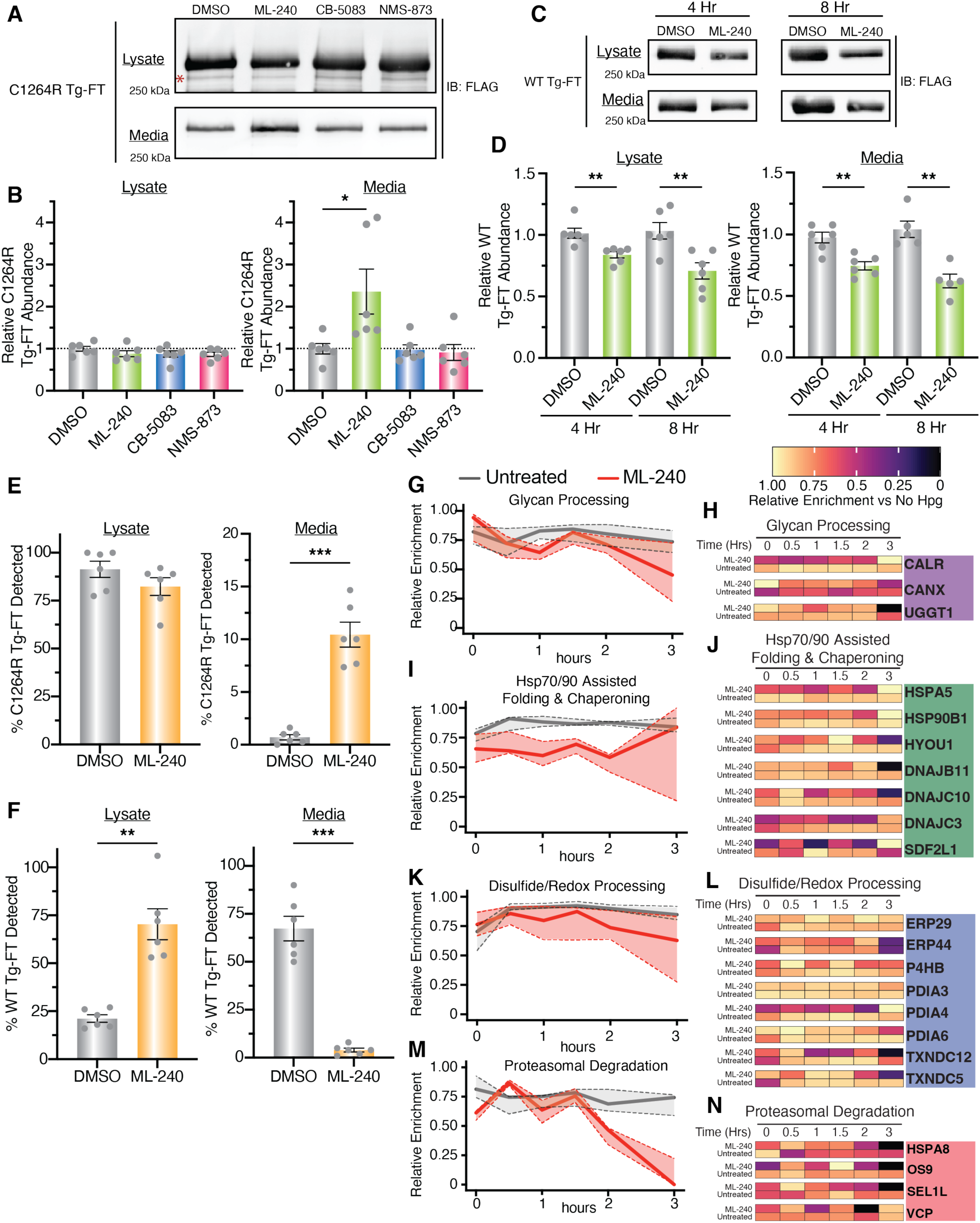
Secretion Rescue of C1264R Tg is Associated with Temporal Remodeling of the Tg Interactome. (A - B) Western blots analysis (A) and quantification (B) of C1264R Tg-FT FTR cells treated with VCP inhibitors ML-240 (10μM), CB-5083 (5μM), NMS-873 (10μM), or vehicle (0.1% DMSO) for 8 h. Lysate and media samples were subjected to immunoprecipitation and analyzed via immunoblotting. Data is normalized to the median C1264R Tg-FT abundance of DMSO treated samples. Data represented as mean ± SEM. Statistical testing performed using an unpaired student’s t-test with Welch’s correction, *p<0.05, **p<0.005, ***p<0.0005. N = 6. (C -D) Western blots analysis (C) and quantification (D) of WT Tg-FT FTR cells treated with ML-240 for 4 and 8 h. Lysate and media samples were processed and analyzed as described above. Data is normalized to the median WT Tg-FT abundance of DMSO treated samples. Data representation and statistical testing as in (B). N = 6. (E) ^35^S Pulse-chase analysis of C1264R Tg-FT FTR cells with ML-240 treatment. Cells were pre-treated with ML-240 or DMSO for 15 minutes prior to pulse labeling with ^35^S for 30 minutes and chased for 4 h with DMSO or ML-240 treatment. Data is represented as mean ± SEM. Full data panel available in Fig EV9A – B. N = 6. (F) ^35^S Pulse chase analysis of WT Tg-FT FRT cells with ML-240 (10μM) treatment. Samples processed as described above in (E). Data is represented as mean ± SEM. Full data panel available in Fig EV9C - D. N = 6. (G-N) TRIP analysis showing the relative enrichment for select Tg interactors across proteostasis pathways between untreated and ML-240 (10μM) treated C1264R Tg. Cells were treated with ML-240 (10μM) and analyzed utilizing the TRIP workflow (Fig 2). The average log2 fold change enrichment values across time points for a given interactor are used to scale data. Positive enrichment values were scaled from 0 to 1. The enrichment is shown for Glycan Processing (G-H), Hsp70/90 Assisted Folding (I-J), Disulfide/Redox Processing (K-L), and Proteasomal Degradation (M-N). Aggregate pathway enrichment is displayed in (G), (I), (J) and (M), where lines represent the median scaled enrichment for the group of interactors and shades correspond to the first and third quartile range. Heatmaps with individual enrichments of interactors is shown in (H), (J), (L), and (N). Data available in **Dataset EV4**.

We hypothesized that ML-240 treatment may differentially regulate Tg degradation and processing in a construct specific manner. Hence, we turned to a ^35^S pulse-chase assay to fully characterize Tg degradation and processing dynamics and found that lysate abundance of C1264R Tg was not significantly changed at the 4 h time point with ML-240 treatment compared to DMSO (**Fig 7E**, **Fig EV9A,B**). This paralleled the previous results with VCP silencing and ML-240 treatment under steady-state conditions in FRT cells. Impressively, C1264R Tg secretion was increased 10-fold at the 4 h time point with ML-240 treatment compared to DMSO with no significant change in C1264R Tg degradation in the presence of ML-240, (**Fig 7E**, **Fig EV9A,B**). Conversely, for WT Tg ML-240 treatment led to a significant increase in lysate accumulation at 4 hcompared to DMSO (**Fig 7F**, **Fig EV9C,D**). This increase in lysate abundance correlated with a significant decrease in WT Tg secretion from 67% to 4% in the presence of ML-240 compared to DMSO, without altering WT degradation (**Fig 7F**, **Fig EV9C,D**).

To understand whether this rescue in secretion was uniquely linked to VCP inhibition or could be more broadly attributed to blocking Tg degradation, we tested the proteasomal inhibitor bortezomib, and lysosomal inhibitor bafilomycin. Bafilomycin increased WT Tg lysate abundance, and bortezomib significantly increased A2234D lysate abundance, consistent with a role of these degradation processes in Tg PQC (**Fig EV10A**). When monitoring Tg-NLuc media abundance, neither bafilomycin nor bortezomib significantly altered WT, A2234D, or C1264R abundance, confirming that general inhibition of proteasomal or lysosomal degradation does with rescue mutant Tg secretion (**Fig EV10B**).

Finally, we monitored activation of the unfolded protein response (UPR) in the presence of ML-240 in FRT cells expressing C1264R Tg-FT. Phosphorylation of eIF2α, an activation marker for the PERK arm of the UPR, was induced within 2 h of ML-240 treatment (**Fig EV10C**). We further investigated the induction of UPR targets via qRT-PCR. HSPA5 and ASNS transcripts, markers of ATF6 and PERK UPR activation respectively, remained unchanged or slightly decreased after 3 h treatment with ML-240 in C1264R Tg cells (**Fig EV10D**). Only DNAJB9, a marker of the IRE1 arm of the UPR, showed a significant increase in both WT Tg and C2164R Tg FRT cells (**Fig EV10D**). Moreover, ML-240 did not significantly alter cell viability after 3 h, as measured by propidium iodide staining (**Fig EV10E**). Overall, these results highlight that the short ML-240 treatment induces early UPR markers, but the selective rescue of C1264R Tg secretion via ML-240 treatment is unlikely the results of global remodeling of the ER PN due to UPR activation.

### Secretion Rescue of C1264R Tg is Associated with Temporal Remodeling of the Interactome

After identifying that VCP inhibition via ML-240 rescued C1264R Tg secretion, we sought to use TRIP to capture PQC interaction changes that correlated with increased secretion. TRIP was carried out in FRT cells expressing C1264R Tg in the presence of ML-240 to monitor temporal changes in PN interactions (**Appendix Fig S3, Dataset EV4**). Both glycan processing and Hsp70/90 chaperoning pathways exhibited broad decreases in C1264R Tg interactions in ML-240 treated samples compared to untreated samples (**Fig 7G,H**). Particularly, CALR, CANX and UGGT1 interactions tapered off more rapidly within 0.5 h compared to untreated C1264R. In contrast, interactions with key Hsp70/90 chaperones components remained relatively steady throughout the chase period before peaking at the 3 h time point (**Fig 7I,J**). Conversely, C1264R interactions with chaperones HSPA5 and HSP90B1, and co-chaperones DNAJB11, DNAJC10, and DNAJC3 decreased and mimicked the WT Tg temporal profile (**Fig 3D**, **Fig 7J**).

Interactions with disulfide/redox processing components exhibited milder but marked declines at intermediate time points with ML-240 treatment compared to untreated samples (**Fig 7K,L**, **Appendix Fig S3**). PDIA4 interactions remained much lower before peaking at the 3 h time point (**Fig 7L**). Conversely, ERP29 and PDIA3 interactions remained largely unchanged in the presence of ML-240 (**Fig 7L**). The most striking interaction changes occurred with proteasomal degradation components, which remained steady until 1.5 h, but then abruptly declined with ML-240 treatment at later time points (**Fig 7M,N**). This decline tracks with changes to the glycan processing machinery, highlighting that the coordination between N-glycosylation and diverting Tg away from ERAD may be a key to the rescue mechanism.

## DISCUSSION

The timing of protein-protein interactions implicated in cellular processes and pathogenic states remains pivotal to our understanding of disease mechanisms. Nonetheless, the temporal measurement of these interactions has remained difficult and proven to be a bottleneck in elucidating the coordination of complex biological pathways such as the PN. Here, we developed an approach to temporally resolve protein-protein interactions implicated in PQC. Furthermore, we complemented these data with a functional genomic screen to further characterize and investigate the implications of these protein-protein interactions on Tg processing. This combination of our novel TRIP approach coupled with functional screening deconvoluted previously established PQC mechanisms for Tg processing while also providing new paradigms within PQC pathways that are critical for secretion of the prohormone.

TRIP has allowed identification and resolution of unique temporal changes in Tg interactions with glycan processing components including CANX, CALR, and UGGT1, while contrasting WT to mutant Tg variants. These changes subsequently correlate with altered interactions with ERAD components EDEM3, OS9, SEL1L, and VCP. Moreover, we identified a broader scope of Tg interactors in thyroid tissue, including ER-phagy receptors ATL3, CCPG1, RTN3, and the RTN3 adaptor protein PGRMC1. Identification of these receptors establishes a direct link from Tg processing to ER-phagy or ERLAD degradation mechanisms. Additionally, glycan dependent and independent mechanisms have been established for the degradation of ERAD resistant ER clients (He *et al*, 2021; Fregno *et al*, 2021). This overlap in degradation pathways may be similarly at play in the case of Tg biogenesis and processing, especially with transient disulfide-linked aggregation taking place during nascent Tg folding and mutants forming difficult to degrade aggregates (Kim *et al*, 1993; di Jeso *et al*, 2005; Menon *et al*, 2007; di Jeso *et al*, 2014).

An important finding was that TRIP was capable of resolving subtle temporal interaction changes, for example with CALR, ERP29, ERP44, P4HB, and VCP, which were otherwise masked in steady-state interactomics data (Wright *et al*, 2021). Nonetheless, there are some limitations within our TRIP methodology. TRIP relies on a two-stage purification strategy which increases sample handling and limits the amount of protein that can be subsequently enriched. Both add inherent variability to the workflow. Furthermore, pulsed labeling with unnatural amino acids has been shown to slow portions of protein translation (Kiick *et al*, 2001; Dieterich *et al*, 2006; Bagert *et al*, 2014; Lang & Chin, 2014). To address this, we utilized a labeling time of 1 h which allows us to generate a large enough labeled population of Tg-FT for TRIP analysis, but some early interactions are likely missed within the TRIP workflow. In the case of mutant Tg, performing the TRIP analysis for much longer chase periods (6-8 h) may provide insightful details to the iterative binding process of PN components that is thought to facilitate protein retention within the secretory pathway. Additionally, within our dataset we noticed that the temporal profiles for ribosomal and proteasomal subunits, trafficking and lysosomal components are inherently difficult to measure across experiments. To efficiently measure these components temporally the TRIP methodology will require further optimization. The functional implications of protein-protein interactions can be difficult to deduce, especially in the case of PQC mechanisms containing several layers of redundancy across stress response pathways, paralogs, and multiple unique proteins sharing similar functions (Wright & Plate, 2021; Bludau & Aebersold, 2020; Karagöz *et al*, 2019; Braakman & Hebert, 2013). This led us to establish a siRNA screening platform to complement our TRIP data and broadly investigate the functional implications of PQC components. With this assay we found the trafficking factor NAPA (α-SNAP) was heavily implicated in WT Tg secretion. Most strikingly, we found that VCP and TEX264 were implicated in C1264R processing, and siRNA silencing of either led to rescue of C1264R secretion. Rescue of mutant Tg trafficking from ER to Golgi, but not secretion, with low-temperature correction had been documented previously (Kim *et al*, 1996).Yet, this is the first study, to our knowledge, that identified a restorative approach to mutant Tg secretion. To expound upon this, our findings that pharmacological VCP inhibition selectively rescues C1264R secretion compared to WT, and mutant Tg was selectively enriched with TEX264 further corroborated the TRIP data establishing that Tg mutants undergo differential interactions with degradation pathways compared to WT Tg. Similar phenomena have been observed for the partitioning and differential interactions of other prohormone proteins, such as proinsulin and proopiomelanocortin, and proteasome resistant polymers of alpha1-antitrypsin Z-variant. RTN3 is implicated in these differential interactions and is shown to be selective for these prohormones compared to other ER-phagy receptors (Chen *et al*, 2021; Cunningham *et al*, 2019; Fregno *et al*, 2018). While A2234D and C1264R Tg were preferentially enriched with TEX264 compared to WT, it remains unclear what other accessory proteins may be necessary for the recognition of TEX264 clients (Chino *et al*, 2019; An *et al*, 2019). Furthermore, TEX264 function in both protein degradation and DNA damage repair further complicates siRNA-based investigations (Fielden et al., 2022). Further investigation is needed to fully elucidate 1) if Tg degradation takes place via ER-phagy and 2) by which mechanisms this targeting is mediated.

As we discovered that pharmacological VCP inhibition with ML-240 can rescue C1264R Tg secretion yet is detrimental for WT Tg processing, it is unclear whether VCP may exhibit distinct functions for WT and mutant Tg PQC. Finally, as ML-240 is shown to block both the proteasomal and autophagic functions of VCP it is unclear which of these pathways may be playing a role in the rescue of C1264R, or detrimental WT processing (Chou *et al*, 2013, 2014).

We used our TRIP method to monitor the changes in interactions associated with the rescue of C1264R Tg with pharmacological VCP inhibition. We found that this rescue is correlated with broad temporal changes in interactions across glycan processing, Hsp70/90 chaperoning and proteasomal degradation pathways, while exhibiting more discrete changes with select disulfide/redox processing components. Mapping these temporal changes in response to pharmacological VCP inhibition and C1264R rescue highlights the capabilities of TRIP to not only resolve protein-protein interactions across disease states but also identify compensatory mechanisms that may take place with drug treatment or other modulating techniques like gene inhibition or activation used to study or treat disease states. Consequently, TRIP should find broad applicability for delineating the proteostasis deficiencies that give rise to diverse protein misfolding diseases and elucidating other cellular interactome dynamics.

## Materials and methods

All unique/stable reagents generated, including plasmids and cell lines are available from the corresponding author with a complete Materials Transfer Agreement.

### Key resources table

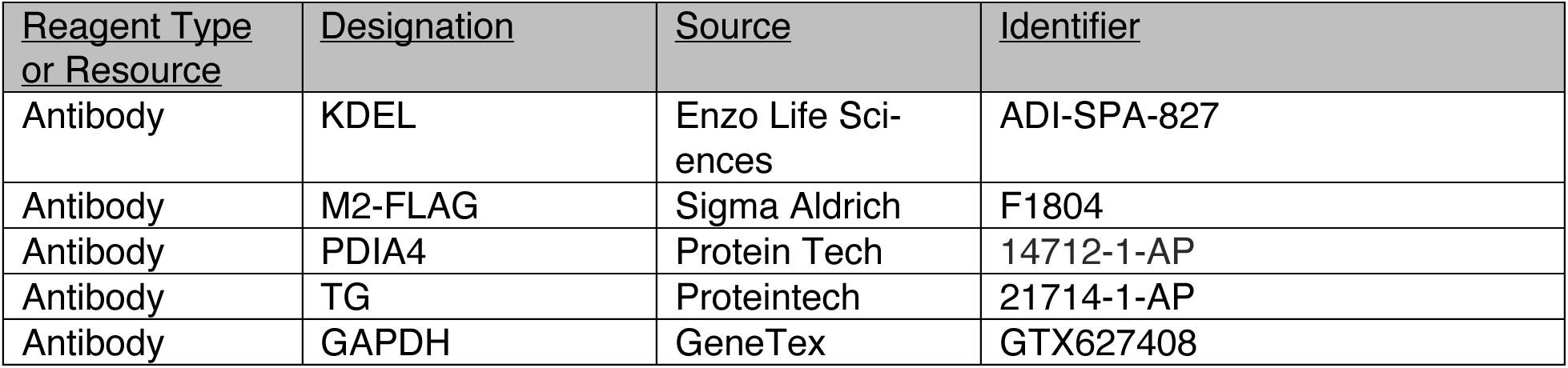
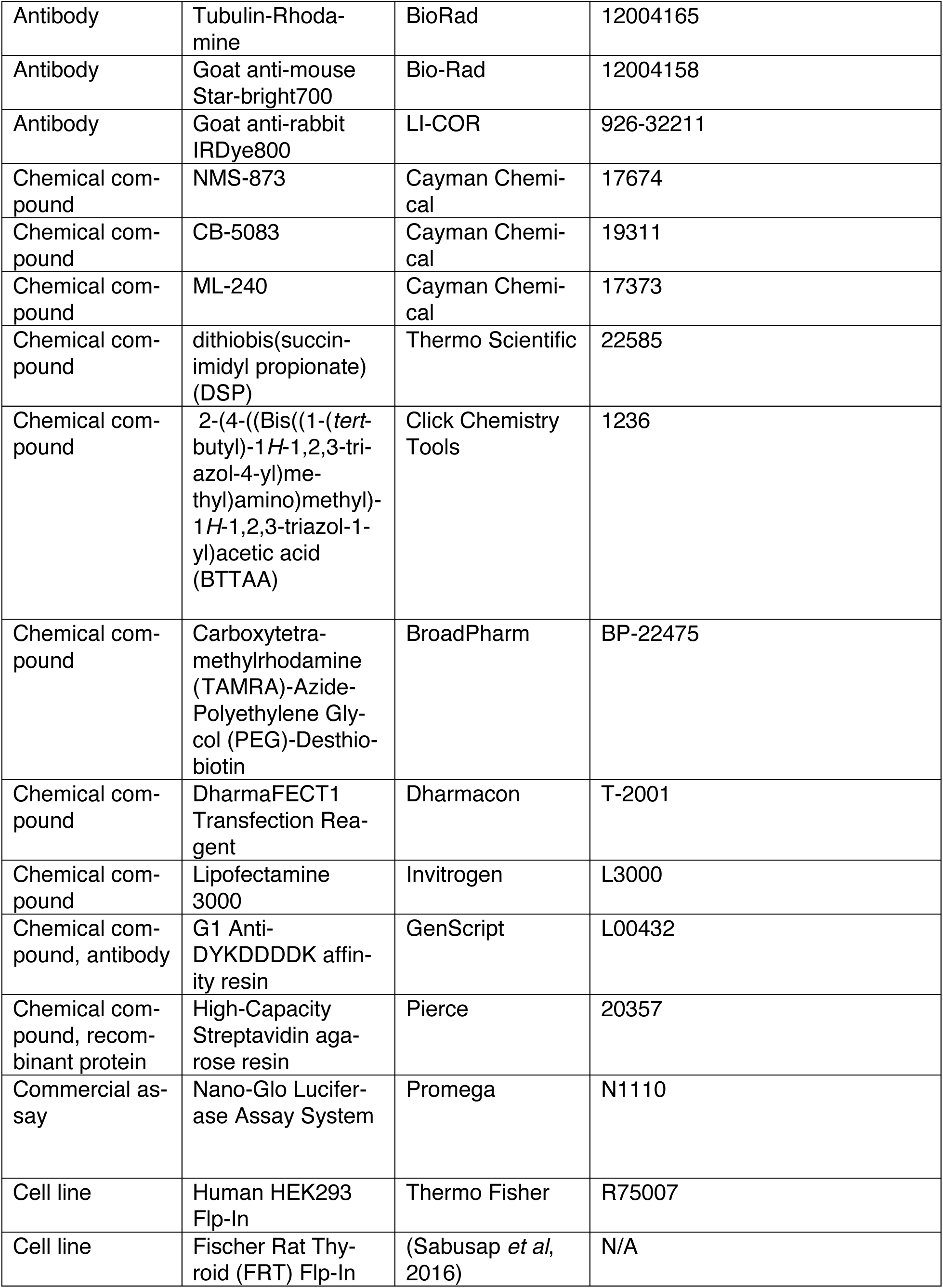
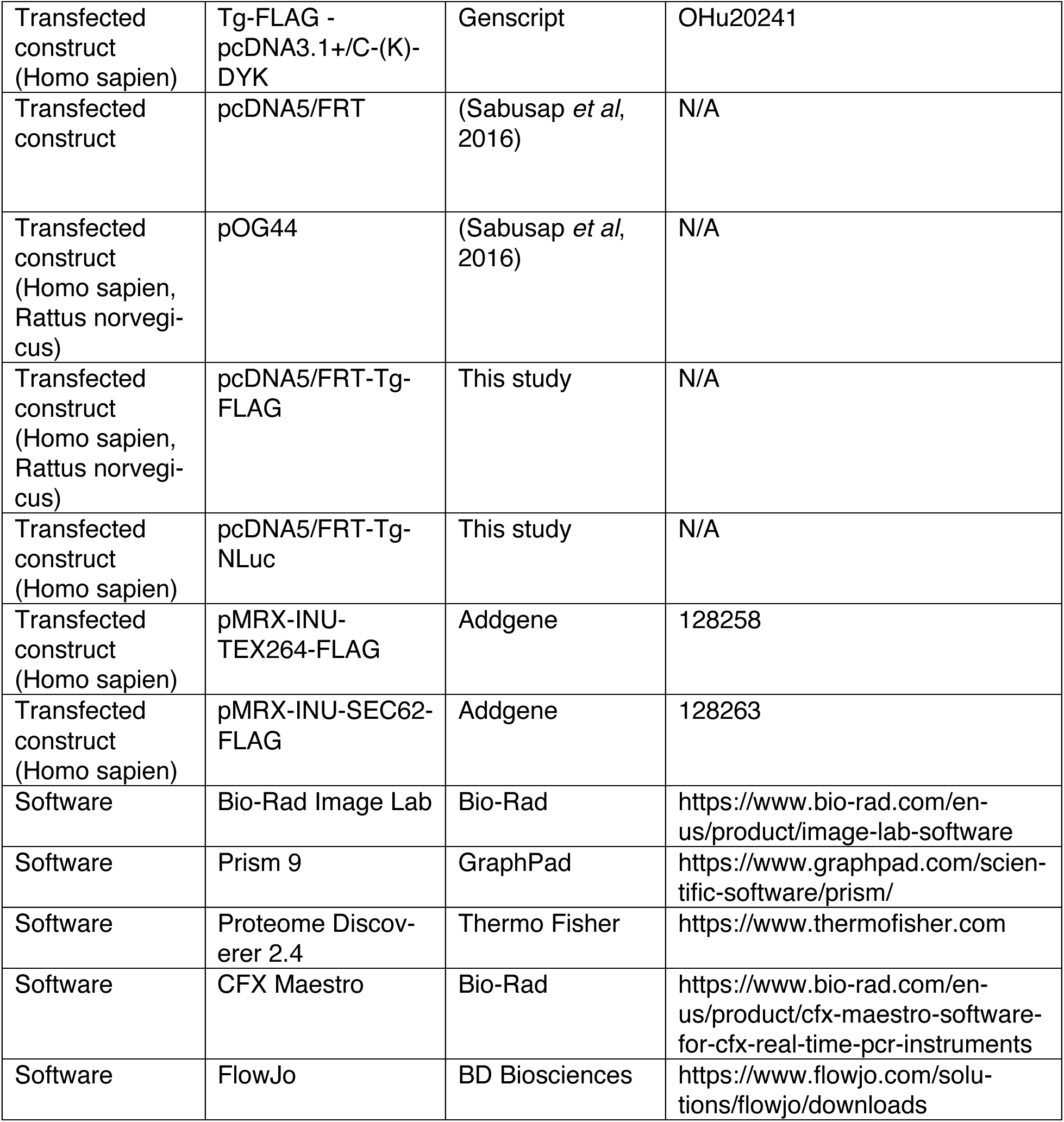

### Plasmid production and antibodies

Tg-FLAG in pcDNA3.1+/C-(K)-DYK plasmid was purchased from Genscript (Clone ID OHu20241). The Tg-FLAG gene was then amplified and assembled with an empty pcDNA5/FRT expression vector using a HiFi DNA assembly kit (New England BioLabs, E2621). This plasmid then underwent site-directed mutagenesis to produce pcDNA5-C1264R-Tg-FLAG, and pcDNA5-A2234D-Tg-FLAG plasmids.

An oligonucleotide fragment encoding Nanoluciferase (NLuc) was ordered from Genewiz. To generate pcDNA5/FRT-Tg-NLuc plasmids the Tg gene was amplified from the pcDNA5/FRT-Tg-FLAG plasmid and assembled with the NLuc fragment using a HiFi DNA assembly kit (New England BioLabs). To generate the respective mutant construct plasmids for A2234D and C1264R Tg, site directed mutagenesis was performed using a Q5 polymerase (New England BioLabs). All oligonucleotide sequences used for site-directed mutagenesis can be found in **Table EV2**.

### Cell line engineering

FRT cells were cultured in Ham’s F-12, Coon’s Modification (F12) media (Sigma, cat. No. F6636) supplemented with 5% fetal bovine serum (FBS), and 1% penicillin (10,000U) / streptomycin (10,000 μg/mL). To generate FRT flp-Tg-FT cells, 5×10^5^ cells cultured for 1 day were cotransfected with 2.25 μg flp recombinase pOG44 plasmid and 0.25 μg of FT-Tg pcDNA5 plasmid using Lipofectamine 3000. Cells were then cultured in media containing Hygromycin B (100 μg/mL) to select site-specific recombinants. Resistant clonal lines were sorted into single cell colonies using flow cytometry and screened for FT-Tg expression (**Fig EV1**).

To generate HEK293 flp-Tg-NLuc cells, 5×10^5^ cells cultured for 1 day were cotransfected with 2.25 μg flp recombinase pOG44 plasmid and 0.25 μg of NLuc-Tg pcDNA5 plasmid using Lipofectamine 3000. Cells were then cultured in media containing Hygromycin B (100 μg/mL) to select site specific recombinants. Polyclonal lines were screened for Tg-NLuc expression and furimazine substrate turnover (**Fig EV6**).

### Time-resolved interactome profiling

Fully confluent 15cm tissue culture plates of FRT cells were used. Cells were washed with phosphate buffered saline (PBS) and depleted of methionine by incubating with methionine free Dulbecco’s Modified Eagle Medium (DMEM) supplemented with 5% dialyzed fetal bovine serum (FBS), 1% L-glutamine (2mM final concentration), 1% L-cysteine (200 μM final concentration), and 1% penicillin (10,000U) / streptomycin (10,000 μg/mL) for 30 minutes. Cells were then pulse labeled with Hpg-enriched DMEM supplemented with with 1% Hpg (200μM final concentration), 5% dialyzed FBS, 1% L-glutamine (200mM final concentration), 1% L-cysteine (200μM), and 1% penicillin (10,000U) / streptomycin (10,000 μg/mL) for 1 h. After pulse labeling cells were washed with F12 media containing 10-fold excess methionine (2mM final concentration). Cells were then cultured in normal F12 media supplemented with 5% FBS and chased for the specified timepoints. Cells were harvested by washing with PBS and then crosslinked with 0.5mM DSP in PBS at 37°C for 10 minutes. Crosslinking was quenched by the addition of 100mM Tris pH 7.5 at 37°C for 5 minutes. Lysates were prepared by lysing in Radioimmunoprecipitation assay (RIPA) buffer (50mM Tris pH 7.5, 150mM NaCl, 0.1% SDS, 1% Triton X-100, 0.5% deoxycholate) with protease inhibitor cocktail (Roche, 4693159001). Protein concentration was normalized to 1mg/mL using a BCA assay (Thermo Scientific, 23225) and cell lysates underwent click reactions with the addition of 0.8mM copper sulfate (diluted from a 20mM stock in water), 1.6mM BTTAA (diluted from a 40mM stock in water), 5mM sodium ascorbate (diluted from a 100mM stock in water), and 100uM TAMRA-Azide-PEG-Desthiobiotin ligand (diluted from a 5mM stock in DMSO). Samples were allowed to react at 37° for 1 h while shaking at 750rpm. Cell lysates were then precleared on 4B sepharose beads (Sigma, 4B200) at 4°C for 1 h while rocking. Precleared lysates were immunoprecipitated with G1 Anti-DYKDDDDK affinity resin overnight at 4°C while rocking. Resin was washed four times with RIPA buffer, and proteins were eluted twice in 100μL immunoprecipitation elution buffer (2% SDS in PBS) by heating at 95°C for 5 minutes. Eluted samples from FLAG immunoprecipitations were then diluted with PBS to reduce the final SDS concentration to 0.2%. The solutions then underwent streptavidin enrichment with high-capacity streptavidin agarose resin (Pierce, 20359) for 2 hat room temperature while rotating. The resin was then washed with 1mL each of 1% SDS, 4M Urea, 1M sodium chloride, followed by a final wash with 1% SDS (all wash buffers dissolved in PBS). Resin was frozen overnight at -80°C and samples were then eluted twice with 100μL biotin elution buffer (50mM Biotin in 1% SDS in PBS) by heating at 37°C and shaking at 750rpm for 1 h. Eluted streptavidin enrichment samples were precipitated in methanol/chloroform, washed three times with methanol, and air dried. Protein pellets were then resuspended in 3μL 1% Rapigest SF Surfactant (Waters, 186002122) followed by the addition of 10μL of 50mM HEPES pH 8.0, and 34.5μL of water. Samples were reduced with 5mM tris(2-carboxyethyl)phosphine (TCEP) (Sigma, 75259) at room temperature for 30 minutes and alkylated with 10mM iodoacetimide (Sigma, I6125) in the dark at room temperature for 30 minutes. Proteins were digested with 0.5 μg of trypsin/Lys-C protease mix (Pierce, A40007) by incubating for 16-18 hat 37°C and shaking at 750rpm. Peptides were reacted with TMTpro 16plex reagents (Thermo Fisher, 44520) in 40% v/v acetonitrile and incubated for 1 h at room temperature. Reactions were quenched by the addition of ammonium bicarbonate (0.4% w/v final concentration) and incubated for 1 h at room temperature. TMT labeled samples were then pooled and acidified with 5% formic acid (Fisher, A117, v/v). Samples were concentrated using a speedvac and resuspended in buffer A (97% water, 2.9% acetonitrile, and 0.1% formic acid, v/v/v). Cleaved Rapigest SF surfactant was removed by centrifugation for 30 minutes at 21,100 x g.

For TRIP analysis coupled with ML-240 treatment, C1264R Tg FRT cells were processed as described above with ML-240 (10μM) supplemented in Hpg-pulse media and throughout the chase period.

### Liquid chromatography – tandem mass spectrometry

MudPIT microcolumns were prepared as previously described (Fonslow *et al*, 2012). Peptide samples were directly loaded onto the columns using a high-pressure chamber. Samples were then desalted for 30 minutes with buffer A (97% water, 2.9% acetonitrile, 0.1% formic acid v/v/v). LC-MS/MS analysis was performed using an Exploris480 (Thermo Fisher) mass spectrometer equipped with an Ultimate3000 RSLCnano system (Thermo Fisher). MudPIT experiments were performed with 10μL sequential injections of 0, 10, 30, 60, and 100% buffer C (500mM ammonium acetate in buffer A), followed by a final injection of 90% buffer C with 10% buffer B (99.9% acetonitrile, 0.1% formic acid v/v) and each step followed by a 140 minute gradient from 4% to 80% B with a flow rate of 500nL/minute on a 20cm fused silica microcapillary column (ID 100 μm) ending with a laser-pulled tip filled with Aqua C18, 3μm, 125 Å resin (Phenomenex). Electrospray ionization (ESI) was performed directly from the analytical column by applying a voltage of 2.2kV with an inlet capillary temperature of 275°C. Data-dependent acquisition of mass spectra was carried out by performing a full scan from 400-1600m/z at a resolution of 120,000. Top-speed data acquisition was used for acquiring MS/MS spectra using a cycle time of 3 seconds, with a normalized collision energy of 32, 0.4m/z isolation window, automatic maximum injection time and 100% normalized AGC target, at a resolution of 45,000 and a defined first mass (m/z) starting at 110. Peptide identification and TMT-based protein quantification was carried out using Proteome Discoverer 2.4. MS/MS spectra were extracted from Thermo Xcalibur .raw file format and searched using SEQUEST against a Uniprot rat proteome database supplemented with the human thyroglobulin gene (accessed 03/2014 and containing 28860 entries). The database was curated to remove redundant protein and splice-isoforms. Searches were carried out using a decoy database of reversed peptide sequences and the following parameters: 20ppm peptide precursor tolerance, 0.02 Da fragment mass tolerance, minimum peptide length of 6 amino acids, trypsin cleavage with a maximum of two missed cleavages, dynamic methionine modification of +15.995 Da (oxidation), dynamic protein N-terminus +42.011 Da (acetylation), -131.040 (methionine loss), -89.030 (methionine loss + acetylation), static cysteine modification of +57.0215 Da (carbamidomethylation), and static peptide N-terminal and lysine modifications of +304.2071 Da (TMTpro 16plex).

### Immunoblotting and SDS-PAGE

Cell lysates were prepared by lysing in RIPA buffer with protease inhibitor cocktail (Roche) and protein concentrations were normalized using a BCA assay (Thermo Scientific). Lysates were then denatured with 1X Laemmli buffer + 100mM DTT and heated at 95°C for 5 minutes before being separated by SDS-PAGE. Samples were transferred onto poly-vinylidene difluoride (PVDF) membranes (Millipore, IPFL00010) for immunoblotting and blocked using 5% non-fat dry milk dissolved in tris buffered saline with 0.1% Tween-20 (Fisher, BP337-100) (TBS-T). Primary antibodies were incubated either at room temperature for 2 h, or overnight at 4°C. Membranes were then washed three times with TBS-T and incubated with secondary antibody in 5% non-fat dry milk dissolved in TBS-T either at room temperature for 1 h or overnight at 4°C. Membranes were washed three times with TBS-T and then imaged using a ChemiDoc MP Imaging System (Bio-Rad). Primary antibodies were acquired from commercial sources and used at the indicated dilutions in immunoblotting buffer (5% bovine serum albumin (BSA) in Tris-buffered saline pH 7.5, 0.1% Tween-20, and 0.1% sodium azide): KDEL (1:1000), M2 anti-FLAG (1:1000), PDIA4 (1:1000), thyroglobulin (1:1000). Tubulin-Rhodamine primary antibody was obtained from commercial sources and used at 1:10000 dilution in 5% milk in Tris-buffered saline pH 7.5, 0.1% Tween-20 (TBS-T). Secondary antibodies were obtained from commercial sources and used at the indicated dilutions in 5% milk in TBS-T: Goat anti-mouse Star-bright700 (1:10000), Goat antirabbit IRDye800 (1:10000).

### qRT-PCR

RNA was prepared from cell pellets using the Quick-RNA miniprep kit (Zymo Research). cDNA was synthesized from 500ng total cellular RNA using random hexamer primer (IDT), oligo-dT primer (IDT), and M-MLV reverse transcriptase (Promega). qPCR analysis was performed using iTaq Universal SYBR Green Supermix (Bio-Rad) combined with primers for genes of interest and reactions were run in 96-well plates on a Bio-Rad CFX qPCR instrument. Data analysis was then carried out in CFX Maestro (Bio-Rad). All oligonucleotide sequences used for qRT-PCR can be found in **Table EV2**.

### siRNA screening assay

Proteins previously identified as Tg interactors, and other key proteostasis network components were selected and targeted using an siGENOME SMARTpool siRNA library (Dharmacon) (Wright *et al*, 2021). HEK293 cells stably expressing Tg-NLuc constructs were seeded into 96-well plates at 2.5×10^4^ cells/well and transfected using DharmaFECT 1 following the DharmaFECT1 protocol (Dharmacon) with a 25nM siRNA concentration. Approximately 36 hafter transfection, cells were washed with PBS and replenished with fresh DMEM media. After 4 h, Tg-NLuc abundance in the lysate and media were measured using the nano-glo luciferase assay system according to the manufactures protocol (Promega). Four controls were included for the experiments including a non-targeting siRNA control, a siGLO fluorescent control to monitor transfection efficiency, a vehicle control containing transfection reagents but lacking any siRNAs, and a lethal TOX control. Data was median normalized across individual 96-well plates (Chung *et al*, 2008). Data represents two independent experiments for WT-NLuc and A2234D-NLuc, and three independent experiments for C1264R NLuc. Cutoff criteria for hits were set to those genes that increased or decreased Tg-NLuc abundance in lysate or media by 3σ.

### FRT siRNA validation studies

For siRNA silencing follow-up experiments, Tg FRT cells were seeded into 6-well dishes at 6.0×10^5^ cells/well transfected using DharmaFECT 1 following the DharmaFECT protocol (Horizon) with a 25nM siRNA concentration. For NAPA complementation experiments, siRNA (25nM) and corresponding plasmid (0.83 µg per 6-well dish) were cotransfected with DharmaFECT Duo (Horizon) Approximately 36 hafter transfection, cells were washed with PBS and harvested for qRT-PCR for Western blotting. For qRT-PCR siRNA target transcript levels were normalized to a GAPDH loading control to monitor siRNA silencing efficiency. For immunoblot analysis approximately 36 hafter transfection cells were washed with PBS and replenished with fresh DMEM media. After 4 hin the case of WT Tg and 8 hin the case of C1264R or A2234D Tg, cells and media samples were harvested. Cells were lysed with 1mL of RIPA with protease inhibitor cocktail (Roche), and lysate and media samples were subjected to immunoprecipitation with G1 Anti-DYKDDDDK affinity resin overnight at 4°C while rocking. After three washes with RIPA buffer, protein samples were eluted with 3× Laemmli buffer with 100 mM DTT heating at 95°C for 5 min. Immunoblot quantification was performed using Image Lab Software (BioRad).

### 35S pulse chase assay

Confluent 6-well dishes of FRT cells (approximately 1×10^6^/well) were metabolically labeled in DMEM depleted of methionine and cysteine and supplemented with EasyTag ^35^S Protein Labeling Mix (Perkin Elmer, NEG772007MC), glutamine, penicillin/streptomycin, and 10% FBS at 37°C for 30 minutes. Afterward, cells were washed twice with F12 media containing 10X methionine and cysteine, followed by a burn off period of 10 minutes in normal F12 media. Cells were then chased for the respective time periods with normal F12 media, lysed with 500uL of RIPA buffer with protease inhibitor cocktail (Roche) and 10 mM DTT. Cell lysates were diluted with 500uL of RIPA buffer with protease inhibitor cocktail (Roche) and subjected to immunoprecipitation with G1 anti-DYKDDDDK affinity resin overnight at 4°C. After three washes with RIPA buffer, protein samples were eluted with 3× Laemmli buffer with 100 mM DTT heating at 95°C for 5 minutes. Eluted samples were then separated by SDS-PAGE, and gels were dried and exposed on a storage phosphor screen. Radioactive band intensity was then measured using a Typhoon Trio Imager (GE Healthcare) and quantified by densitometry in Image Lab (BioRad).

For pulse chase analysis coupled with ML-240 treatment, cells were pre-treated with vehicle (0.1% DMSO) or ML-240 (10μM) for 15 minutes prior to ^35^S pulse labeling. Vehicle (0.1% DMSO) or ML-240 (10μM) treatment was then maintained throughout the pulse labeling and chase period.

### VCP pharmacological inhibition studies

For follow-up experiment with VCP inhibitors, confluent 6-well dishes of C1264R Tg-FT FRT cells were washed with PBS and fresh F12 media supplemented with ML-240 (10μM), CB-5083 (5μM), NMS-873 (10μM), or vehicle (0.1% DMSO) was added and incubated for 8 h. For WT Tg-FT cells, confluent 6-well dishes were similarly used, washed with PBS and fresh F12 media supplemented with ML-240 (10μM) or vehicle (0.1% DMSO) was added and incubated for 4 or 8 h. Cells were lysed with 1mL of RIPA with protease inhibitor cocktail (Roche), and lysate and media samples were subjected to immunoprecipitation with G1 Anti-DYKDDDDK affinity resin overnight at 4°C while rocking. After three washes with RIPA buffer, protein samples were eluted with 3× Laemmli buffer with 100 mM DTT heating at 95°C for 5 minutes. Immunoblot quantification was performed using Image Lab Software (BioRad).

Viability with ML-240 was monitored using propidium iodide staining. Briefly, cells were treated with either ML-240 (10μM) or vehicle (0.1% DMSO) for 4 hbefore being harvested and stained with propidium iodide (1μg/mL) at room temperature for 15 minutes in the dark. Cells permeabilized with 0.2% Triton were used as a positive staining control. Data analysis was then carried out in FlowJo (BD Biosciences).

### TEX264 & Tg-NLuc co-immunoprecipitation studies

HEK293 flp-Tg-NLuc cells were cultured at 1.0×10^5^ cells/well in 12-well tissue culture dishes for 1 day and transfected with either a fluorescent control, c-terminal FLAG-tag TEX264, or c-terminal FLAG-tag SEC62 using a calcium phosphate method. Confluent plates were harvested by lysing with 300uL of TNI buffer (50mM Tris pH 7.5, 150 mM NaCl, 0.5% IGEPAL CA-630 (Sigma-Aldrich)) and protease inhibitor (Roche). Lysates were sonicated for 10 minutes at room temperature and normalized using a BCA Assay (Thermo Scientific). Cell lysates were then precleared on 4B sepharose beads (Sigma, 4B200) at 4°C for 1 h while rocking, then immunoprecipitated with G1 Anti-DYKDDDDK affinity resin overnight at 4°C while rocking. Resin was washed four times with TNI buffer and resuspended in 250 μL of TNI buffer. 50 μL aliquots were taken and measured using the nano-glo luciferase assay system according to the manufactures protocol (Promega). For differential enrichment, statistical analysis was performed in Prism 9 (GraphPad) using a one-way ANOVA with post hoc Tukey’s multiple testing corrections.

For western blot analysis HEK293 flp-Tg-NLuc 1.0×10^6^ cells/dish in 10 cm tissue culture dishes for 1 day and transfected with either a fluorescent control, TEX264, or SEC62 using a calcium phosphate method. Cells were collected and lysed using 300 μL of TNI buffer with protease inhibitor (Roche) sonicated at room temperature for 10 minutes. Lysates were normalized, precleared, and immunoprecpitated as described above. Proteins were eluted using 6X Laemmli buffer + 100mM DTT, separated by SDS-PAGE and transferred to PVDF membrane (Millipore) for immunoblotting.

### Mass spectrometry interactomics and TMT quantification data analysis

To identify Tg interactors (-) biotin pulldown vs (-) Hpg samples were processed using the DEP pipeline (Zhang *et al*, 2018). Enriched proteins were determined based on those with a log2 fold change of 2σ and Benjamini-Hochberg adjusted p-value (false discovery rate) of 0.05. For pathway enrichment analysis of identified proteins, EnrichR was used and GO Cellular Component and Molecular Function 2018 terms were used to differentiate secretory pathway associated proteins from background (Chen *et al*, 2013). For time resolved analysis, data were processed in R with custom scripts. Briefly, TMT abundances across chase samples were normalized to Tg TMT abundance as described previously and compared to (-) Hpg samples for enrichment analysis (Wright *et al*, 2021). For relative enrichment analysis, the means of log2 interaction differences were scaled to values from 0 to 1, where a value of 1 represented the time point at which the enrichment reached the maximum, and 0 represented the background intensity in the (-) Hpg channel. Negative log2 enrichment values were set to 0 as the enrichment fell below the background.

Inconsistencies in the quantification of Tg bait protein were observed for replicate 5 of WT Tg Hpg-chase samples, likely due to sample loss during the enrichment. Hence, this replicate was excluded from further time-resolved analysis. All analysis scripts are available as described in the Data and Code Availability section.

## DATA AVAILABILITY

Mass spectrometry spectrum and result files are available via ProteomeXchange under identifier PXD035681. All raw and processed TMT quantification data has been provided (**Datasets EV1-4**). Code used for data analysis and generation of figures is available at github.com/wrighmt1/2022_TRIP.

## Article and author information

### Author Details

**Madison T. Wright**

Department of Chemistry, Vanderbilt University, Nashville, TN, USA

**Contribution:** Conceptualization, Funding acquisition, Data curation, Formal analysis, Visualization, Writing – original draft, Writing – review and editing

**Competing Interest:** No competing interest declared

**Bibek Timalsina**

Department of Chemistry, Vanderbilt University, Nashville, TN, USA

Department of Biological Sciences, Vanderbilt University, Nashville, TN, USA

**Contribution:** Data curation, Formal analysis, Writing – review and editing

**Competing Interest:** No competing interest declared

**Valeria Garcia Lopez**

Department of Biological Sciences, Vanderbilt University, Nashville, TN, USA

**Contribution:** Data curation, Formal analysis, Visualization, Writing – review and editing

**Competing Interest:** No competing interest declared

**Jake Hermanson**

Department of Biological Sciences, Vanderbilt University, Nashville, TN, USA

**Contribution:** Data curation, Formal analysis, Visualization, Writing – review and editing

**Competing Interest:** No competing interest declared

**Sarah Garcia**

Department of Chemistry, Vanderbilt University, Nashville, TN, USA

**Contribution:** Data curation, Formal analysis, Visualization, Writing – review and editing

**Competing Interest:** No competing interest declared

**Lars Plate**

Department of Chemistry, Vanderbilt University, Nashville, TN, USA Department of Biological Sciences, Vanderbilt University, Nashville, TN, USA

**Contribution:** Conceptualization, Funding acquisition, Supervision, Data curation, Formal analysis, Writing – original draft, Writing – review and editing

## Funding

**National Science Foundation – Graduate Research Fellowship Program**

- Madison T. Wright

**National Institute of General Medical Sciences (1R35GM133552)**

- Lars Plate

This content is solely the responsibility of the authors and does not necessarily represent the official views of the National Science Foundation or National Institutes of General Medical Sciences.

## Expanded View Figures

**Figure EV1.**
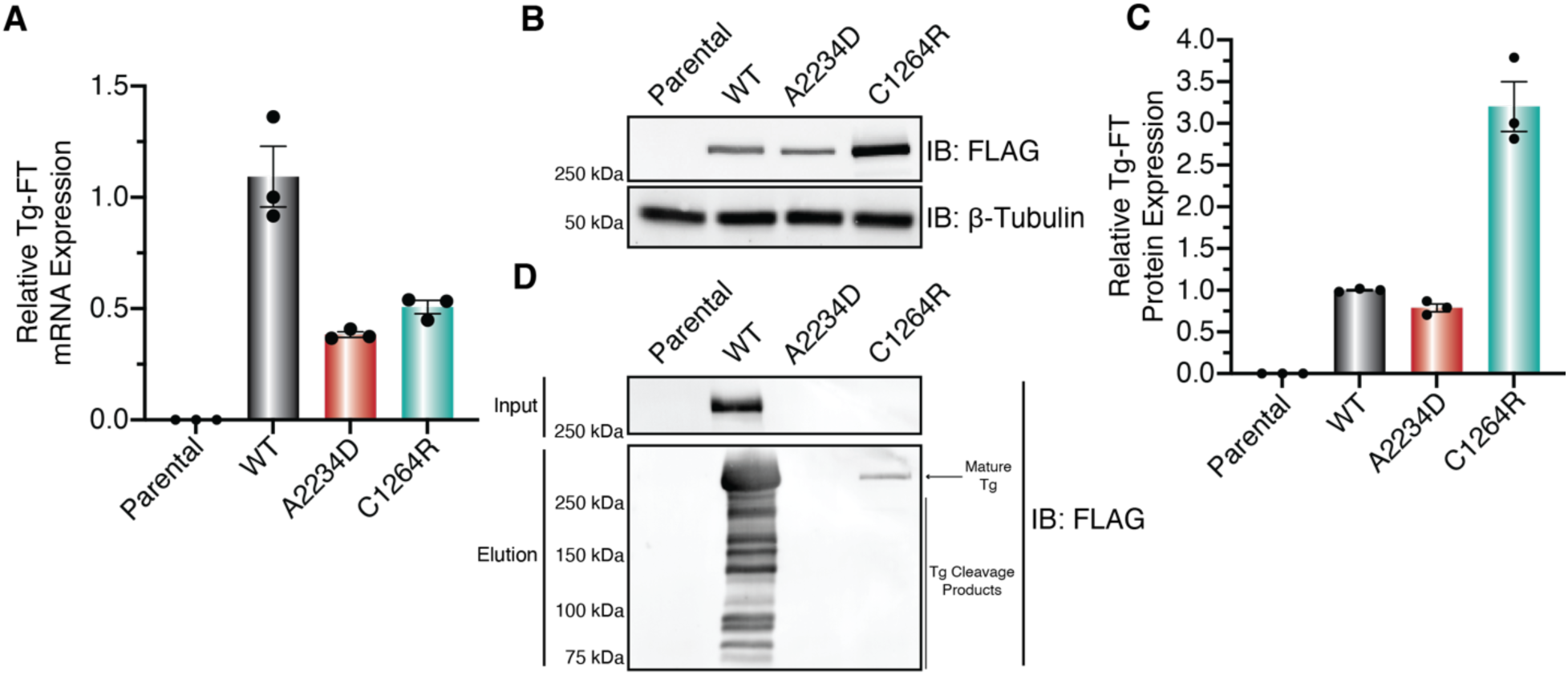
Validation of FRT stable cell lines. (A) Relative expression of Tg-FT RNA from engineered isogenic FRT cells measured by qRT-PCR. After transfections with Tg-FT pcDNA and flp recombinase pOG44, cells were placed under selection with Hygromycin B (100 μg/mL) to select site-specific recombinants. Resistant clonal lines were sorted into single cell colonies using flow cytometry and screen for Tg-FT expression. Data was first normalized to a GAPDH loading control followed by normalization to median WT Tg-FT expression and represented as mean ± SEM. Primers for detection described in **Table EV2**. (B) Western blot analysis of Tg-FT expression in lysates from engineered isogenic FRT cells. FT signal detected via M2-FLAG antibody. FT is only detectable in isogenic cells co-transfected with Tg-FT pcDNA and flp recombinase pOG44, while FT signal is absent in parental cells. β-Tubulin used as a loading control. (C) Quantification of relative Tg-FT expression in engineered FRT cells measured by Western blot analysis in (B). Data was first normalized to the β-Tubulin used as a loading control, followed by normalization to median WT Tg-FT expression and represented as mean ± SEM. (D) Western blot analysis of Tg-FT expression in media from engineered FRT cells. WT Tg-FT is efficiently secreted and detectable in both media inputs and after immunoprecipitation. C1264R secretion is drastically decreased compared to WT Tg-FT and is only detectable after immunoprecipitation, while A2234D secretion is not detectable in inputs or after immunoprecipitation. Mature Tg and Tg cleavage products are annotated in elution samples.

**Figure EV2.**
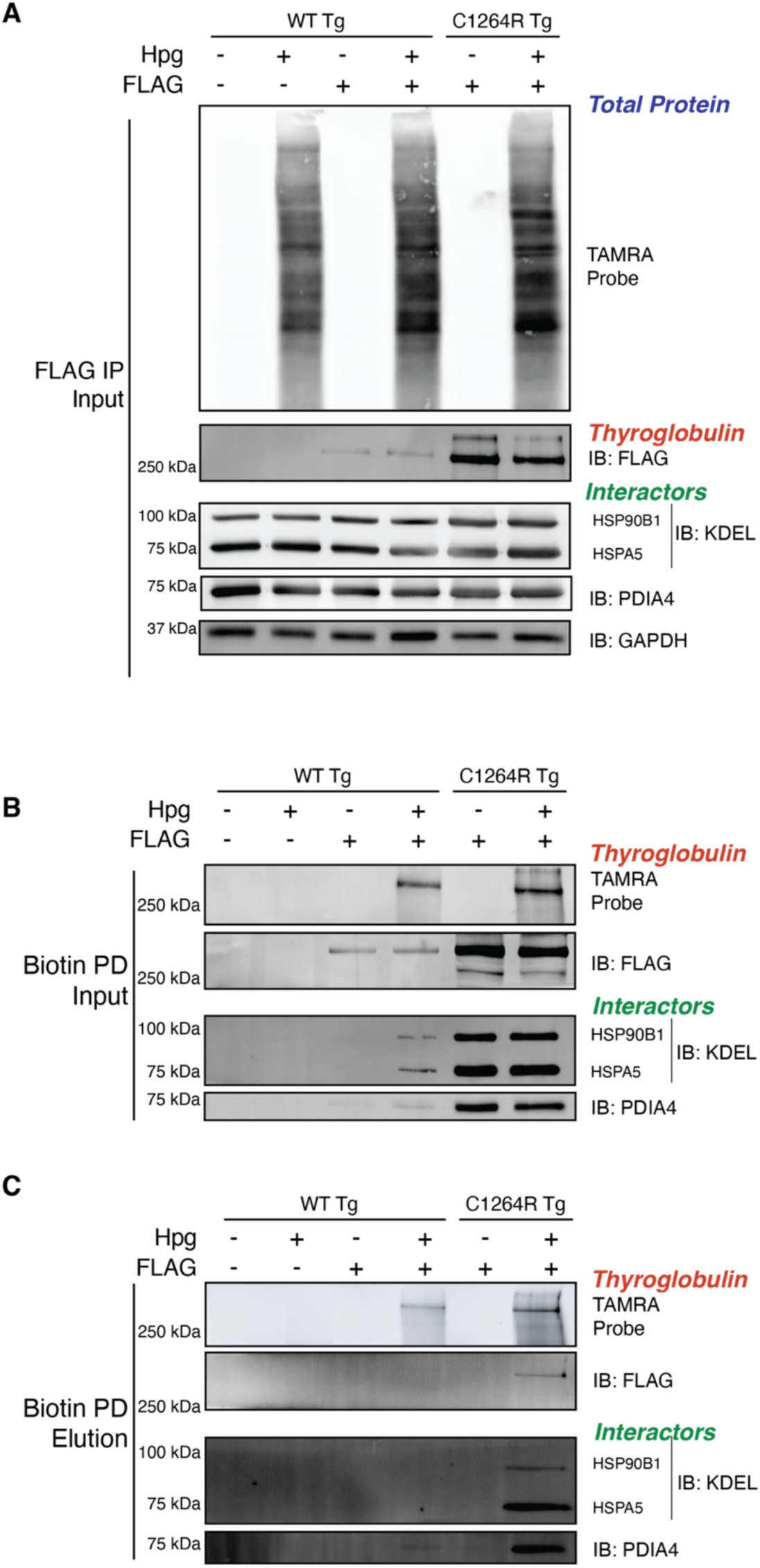
Two-stage Enrichment Strategy Requires FLAG-tag Tg and Pulse-labeling with Hpg. (A-C) Western blot analysis of Tg purification after 4 h of continuous Hpg labeling and functionalization with TAMRA-Azide-PEG-Desthiobiotin probe. All conditions were crosslinked with DSP (0.5mM) for 10 minutes to capture transient interactions. (A) FLAG IP inputs showing TAMRA labeled proteins and immunoblots of Tg (IB: FLAG), interactors, and loading control (IB: GAPDH). (B) FLAG IP elutions showing TAMRA labeled proteins and immunoblots of Tg and interactors. Samples subsequently underwent biotin pulldown. (C) Biotin pulldown elutions with TAMRA labeled proteins, Tg, and interactors.

**Figure EV3.**
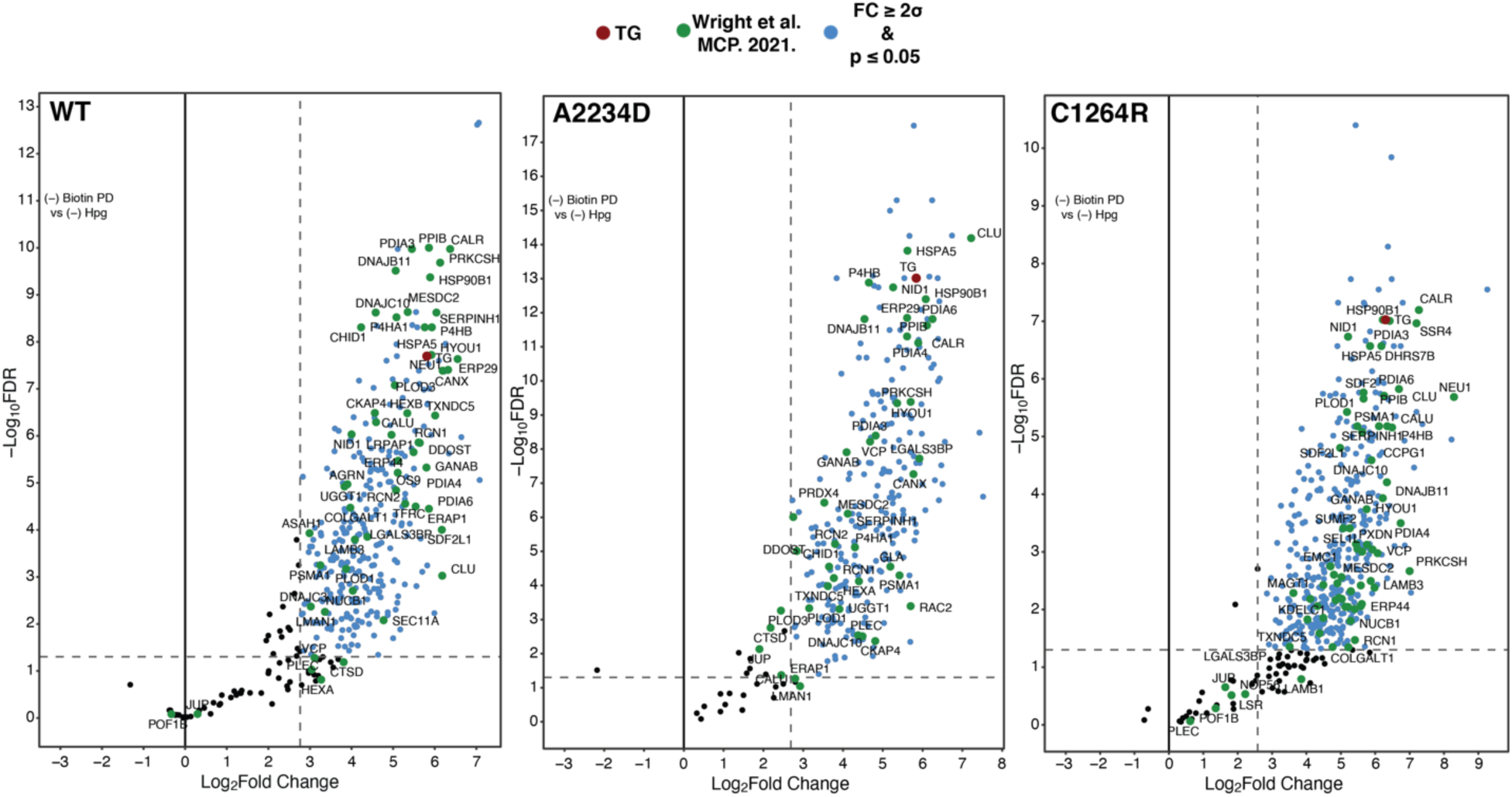
(-) Biotin pulldown carrier samples allow for identification of Tg interactors. Volcano plots for the comparison of (-) biotin pulldown vs (-) Hpg samples to identify Tg interactors in FRT cells. Plots show the average log2 difference vs FDR estimation (Benjamini-Hochberg). Data was processed through the DEP pipeline (script available at github.com/wrighmt1/2022_TRIP) (Zhang et al., 2018). (-) biotin pulldown samples were pulse-labeled with Hpg (200μM) for 1 hour and cultured in normal F12 media throughout the remainder of the 3-hour chase period. Cells were cross linked with DSP (0.5mM) for 10 minutes to capture transient proteoastasis network interactions. Lysates were functionalized with TAMRA-Azide-PEG-Desthiobiotin probe using CuAAC Click reaction. (-) biotin pulldown samples then underwent immunoprecipitation and were processed for mass spectrometry. (-) Hpg samples were processed through the entire dual affinity purification TRIP workflow including 3 hour chase period, absent Hpg labeling, and used for enrichment analysis. Enriched proteins were determined based on those with a log2 fold change of 20 and Benjamini-Hochberg adjusted p-value (false discovery rate) of 0.05. Dashed lines indicate cutoffs for log2 fold change and adjusted p-values. Proteins annotated in blue are above log2 fold change and adjusted p-value cutoffs and considered Tg interactors. Proteins annotated in green were identified as Tg interactors in our previous mass spectrometry dataset (Wright et al., 2021). Tg is annotated in red. Previously identified Tg interactors are heavily enriched in the dataset along with several novel interactors. Full MS data can be found **Dataset EV1** and DEP output can be found in **Dataset EV2**.

**Figure EV4.**
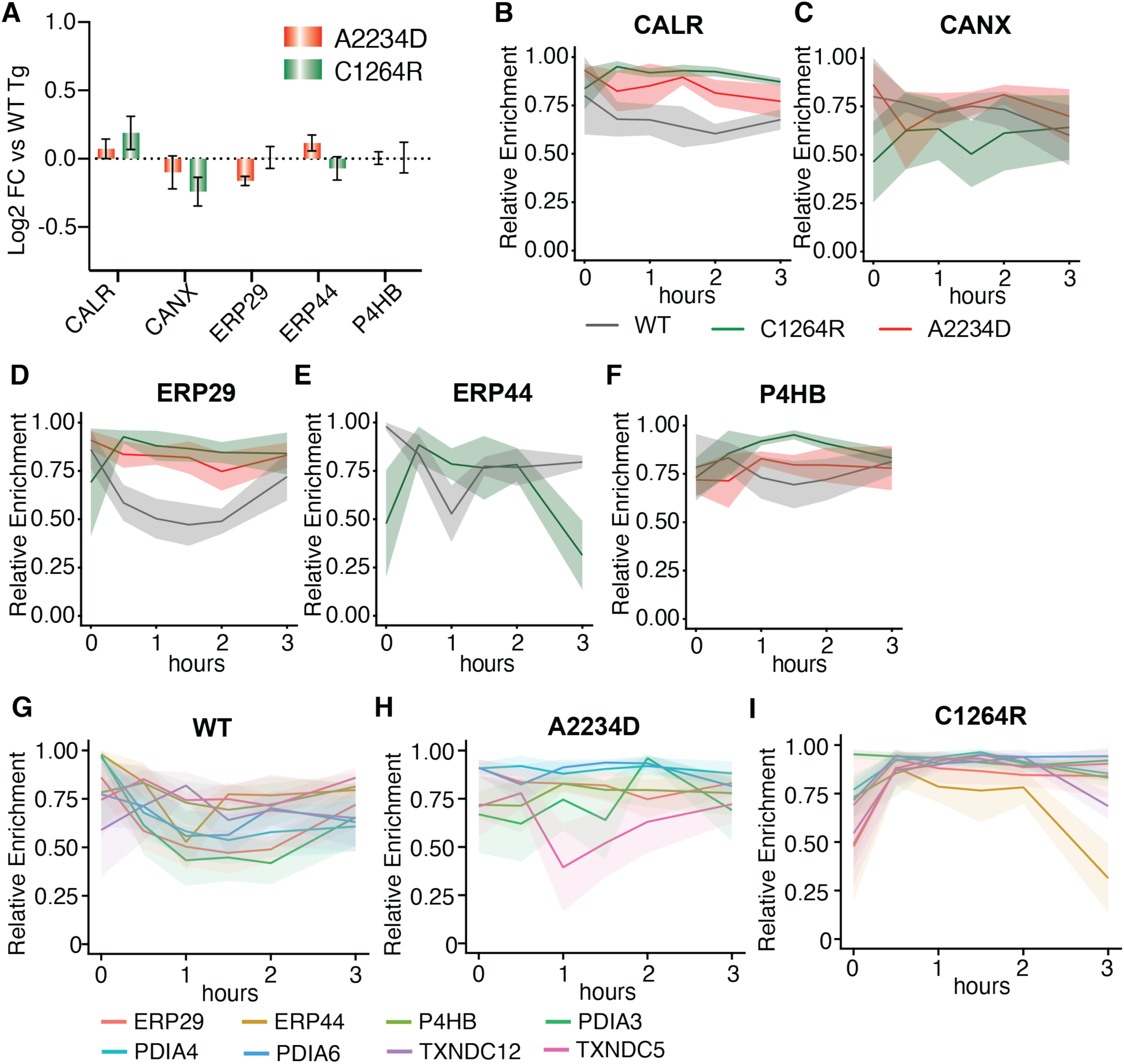
TRIP data for individual interactors. (A) Aggregate (steady-state) interactomics data (right) comparing the enrichment of Tg interactors for mutant Tg to WT Tg (data from Wright et al., 2021). Interactions are mostly unchanged for mutant Tg relative to WT Tg. (B – F) Plots comparing the relative enrichment of interactors calreticulin (CALR, B), calnexin (CANX, C), ERP29 (D), ERP44 (E), and P4HB/PDIA1 (F) throughout the TRIP time course for WT, A2234D, and C1264R Tg. TRIP data can resolve dynamic interaction changes for several mutant Tg interactors, while these changes are muted in the aggregate data. Solid line corresponds to mean and shading represents the SEM (N = 5 for WT Tg; N = 6 for A2234D and C1264R Tg). (G – I) Plots comparing the relative enrichment of individual disulfide/redox processing interactors throughout the TRIP time course for WT (G), A2234D (H), and C1234R (I). Individual protein disulfide isomerases exhibit distinct peak times when interactions reach maximum, thereby revealing an order to their engagement. For instance, PDIA3, PDIA4, PDIA6, and P4HB peak at 0 h for WT Tg, while TXNDC12 peaks later at 1 h. Moreover, the exact temporal sequence of PDI engagements is shifted for A2234D (H) and C1234R Tg (I). Solid line corresponds to mean and shading represents the SEM (N = 5 for WT Tg; N = 6 for A2234D and C1264R Tg). Data available in **Dataset EV4**.

**Figure EV5.**
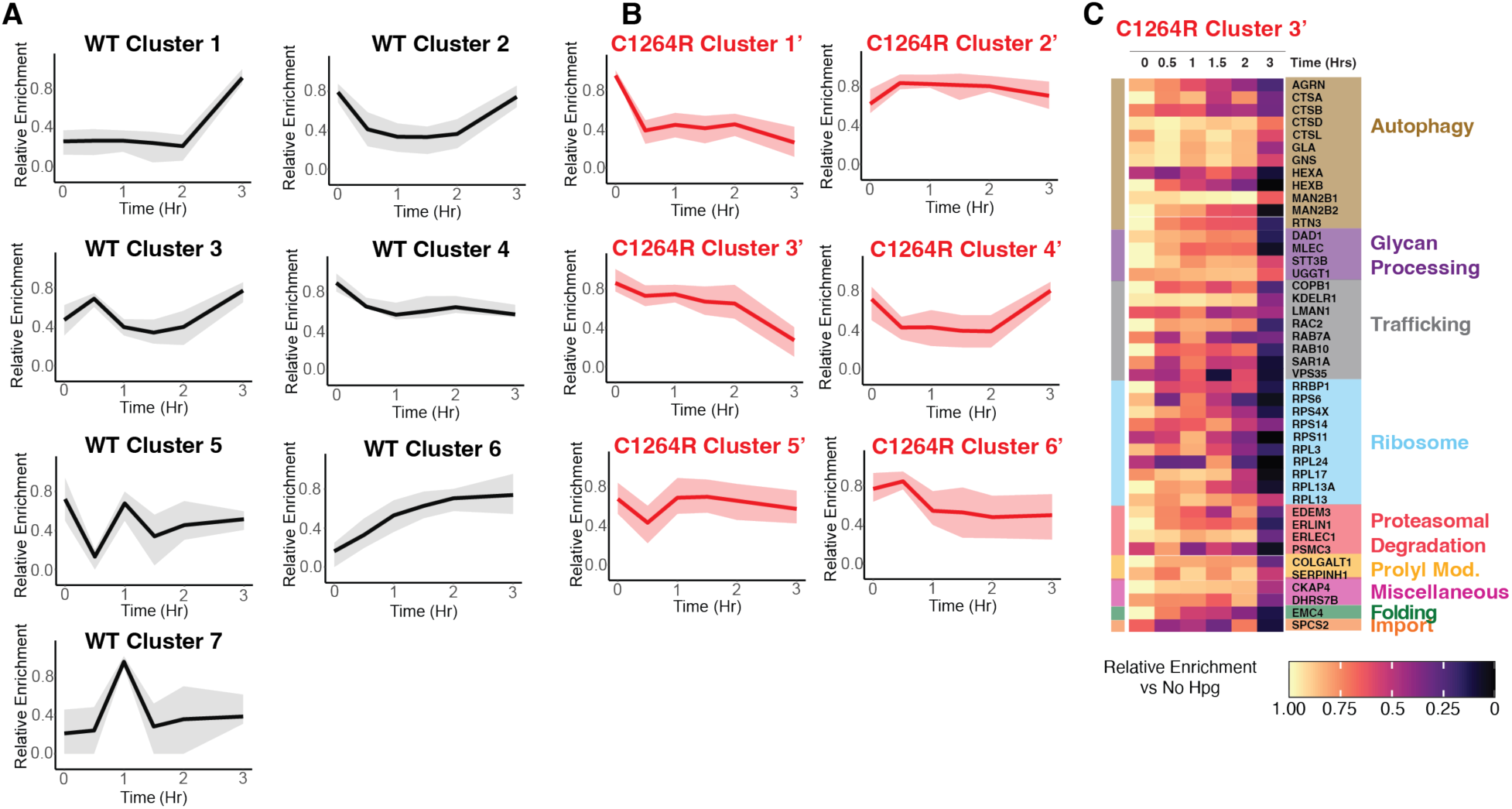
Unbiased clustering of TRIP data. (A – B) Unbiased k-means clustering of TRIP profiles for WT (A) and C1264R (B) to determine co-regulated groups of interactors. k-means clustering was carried out by using the kmeans function from the tslearn python package with the data being normalized using the scaler mean variance function. This analysis resulted in 7 distinct clusters for WT and 6 clusters for C1264R. The line corresponds to the mean scaled log2 fold enrichment and the shading represents the 25 – 75 % quarter range within each cluster. (C) Heatmap for interactors in C1264R Cluster 3, which displayed the strongest interactions at the initial 0 h timepoint. The scaled log2 fold change enrichment for individual interactors is shown, and the individual interactors are grouped by pathway. Several interactors related to autophagy (brown) and glycan processing (purple), including the glycoprotein folding sensor UGGT1, are present in this cluster.

**Figure EV6.**
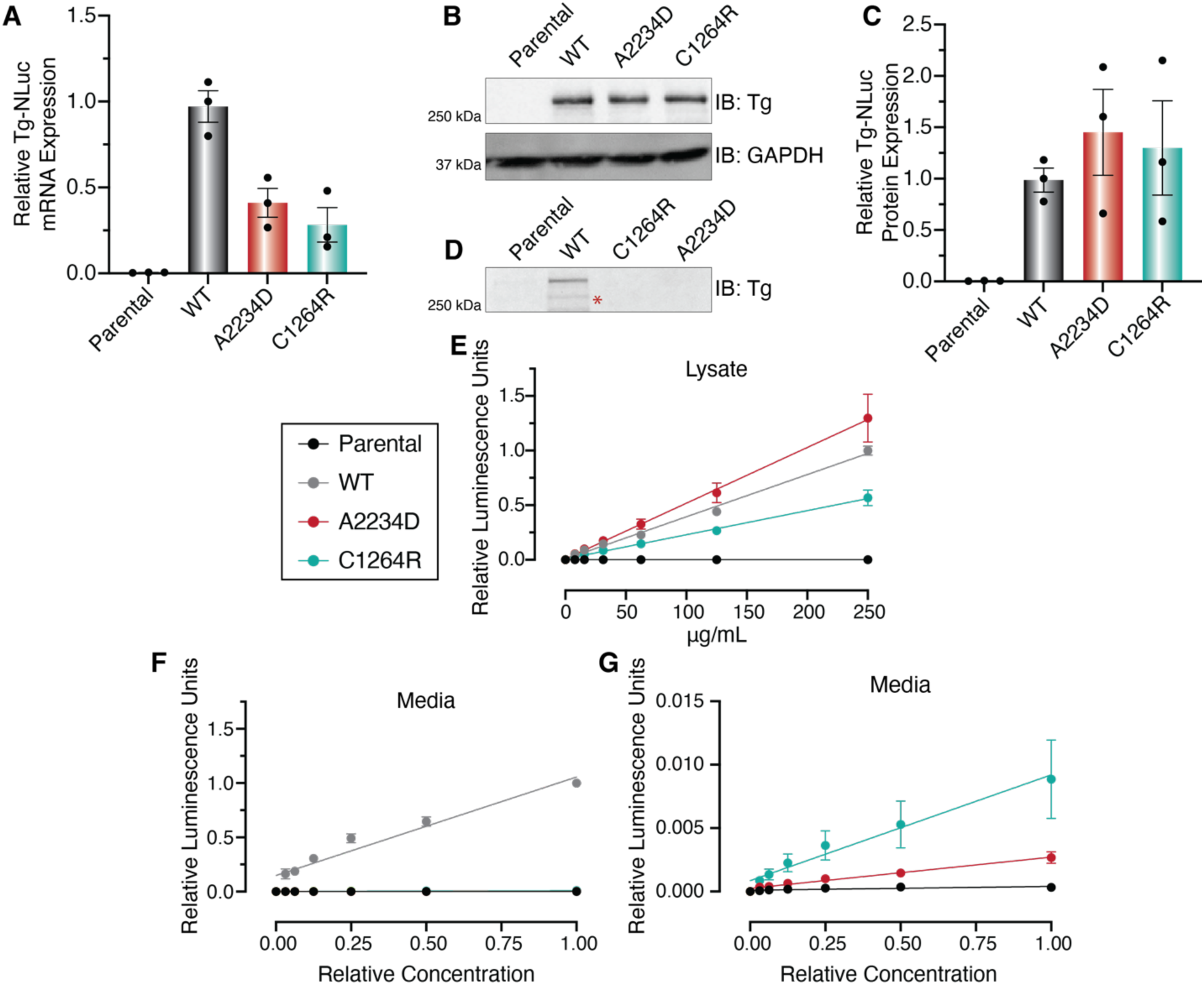
Validation of Tg-NLuc stable cell lines. (A) Relative expression of Tg-NLuc RNA in engineered HEK293 cells measured by qRT-PCR. After transfections with Tg-FT pcDNA and flp recombinase pOG44, cells were placed under selection with Hygromycin B (100 μg/mL) to select site-specific recombinants. Resistant clonal lines were then screened for Tg-NLuc expression. Data was first normalized to a GAPDH loading control followed by normalization to median WT Tg-NLuc expression and represented as mean ± SEM. Primers for detection described in **Table EV2**. (B) Western blot analysis of Tg-NLuc expression in engineered HEK293 cells in lysates. Tg-NLuc signal detected via Tg antibody. Tg is only detectable in isogenic cells co-transfected with Tg-NLuc pcDNA and flp recombinase pOG44, while Tg signal is absent in parental cells. GAPDH used as a loading control. (C) Quantification of relative expression of Tg-NLuc in engineered HEK293 cells measured by western blot analysis in (B). Data first normalized to GAPDH used as a loading control, followed by normalization to median WT Tg-NLuc expression and represented as mean ± SEM. (D) Western blot analysis of Tg-NLuc expression in media from engineered HEK293 cells. WT Tg-NLuc is efficiently secreted and detectable, while C1264R and A2234D secretion is drastically decreased compared to WT Tg-NLuc and is not detectable via western blot analysis. (E-F) Standard curve exhibiting linearity of Tg-NLuc luminescence response based on total protein concentration of lysate (E) and media (F). Lysate or media from Tg-NLuc expressing cells underwent serial dilutions prior to luminescence being measured with the nano-glo luciferase assay system (Promega, N1110). All samples exhibit a linear luminescence signal in response to substrate turnover, while luminescence signal is absent in parental cells. Data is normalized to median WT Tg-NLuc luminescence and represented as mean ± SEM. (G) Rescaled standard curve from (F) exhibiting linearity of mutant Tg-NLuc luminescence response based on relative protein concentration in media. C1264R and A2234D secretion is drastically decreased compared to WT Tg-NLuc but still detectable above background (parental cells). Data is normalized to median WT Tg-NLuc luminescence and represented as mean ± SEM.

**Figure EV7.**
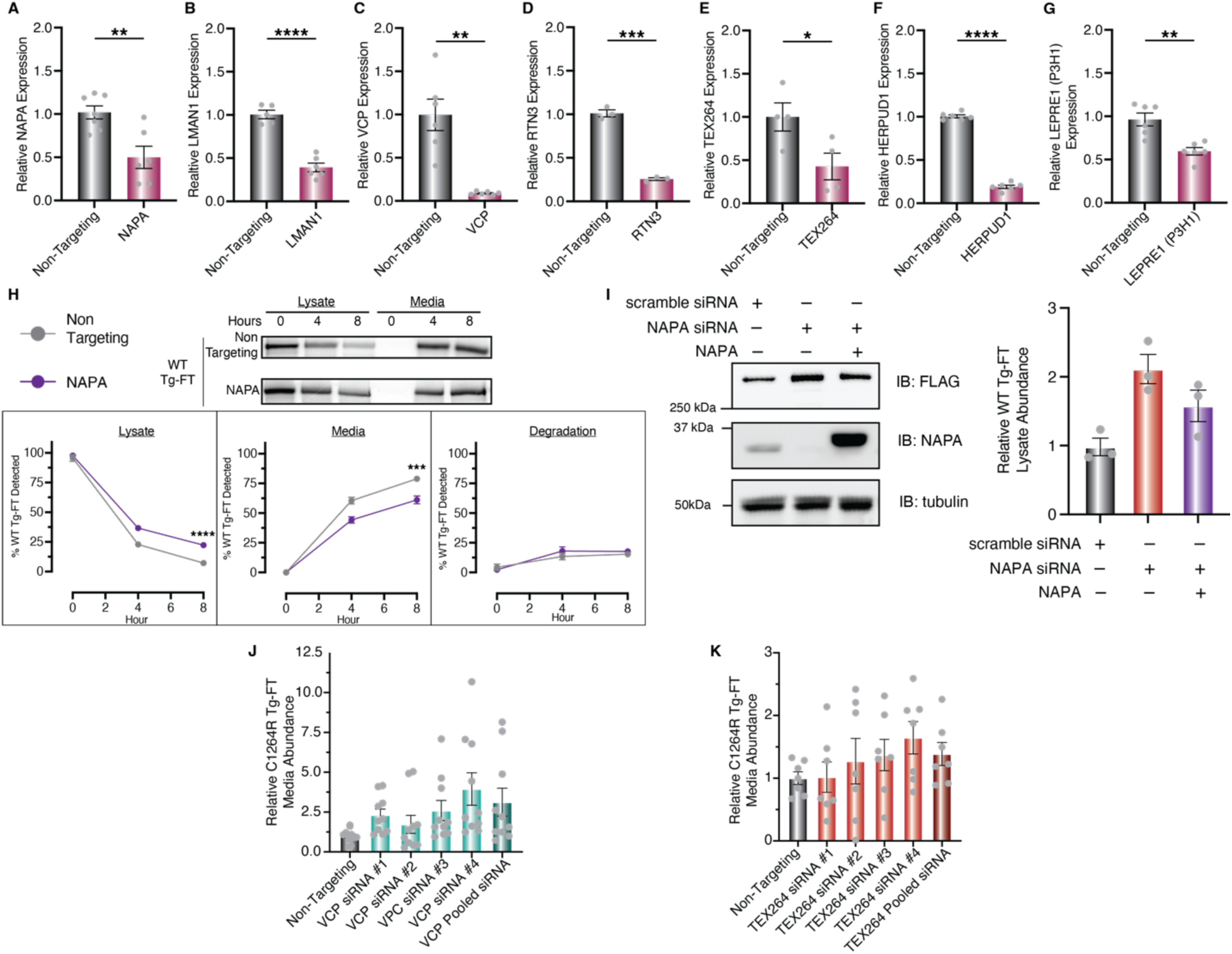
Validation of siRNA screening hits in FRT cells. (A-G) Relative expression of knockdown targets in engineered WT Tg-FT FRT cells with and without siRNA silencing measured by qRT-PCR. Data was first normalized to a GAPDH loading control followed by normalization to median expression of non-targeting transfected samples and represented as mean ± SEM. (A) NAPA (α-SNAP) (B) LMAN1 (C) VCP (D) RTN3 (E) TEX264 (F) HERPUD1 (G) LEPRE11 (P3H1). Statistical testing performed using an unpaired student’s t-test with Welch’s correction, *p<0.05, **p<0.005, ***p<0.0005. Primers for detection described in **Table EV2**. (H) Pulse-chase analysis of WT Tg-FT in FRT cells with NAPA (α-SNAP) siRNA knockdown. Approximately 36 hafter transfection with 25nM siRNAs cells were pulse labeled with EasyTag ^35^S Protein Labeling Mix (Perkin Elmer, NEG772007MC) for 30 minutes and chased for 8 h, collecting samples at 0-, 4-, and 8-h timepoints. Data is normalized to timepoint of maximum Tg recovery and represented as mean ± SEM. % Degradation is defined as 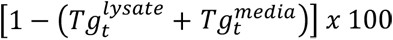 is the fraction of Tg-FT detected in the lysate at a given timepoint *n*, and 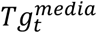 is the fraction of Tg-FT detected in the media at a given timepoint n. Statistical testing performed using an unpaired student’s t-test with Welch’s correction, *p<0.05, **p<0.005, ***p<0.0005. N = 6. (I) Complementation of NAPA knockdown partially reverses WT-Tg retention. FRT cells stably expressing WT-Tg were co-transfected with NAPA siRNA and siRNA resistant NAPA expression plasmid. Cells were harvested 40 h post transfection and lysates were analyzed by Western blot to monitor changes in WT-Tg amounts. Quantification is shown on the right (N = 3). (J-K) Individual siRNA knockdown of VCP (J) and TEX265 (K) in C1264R Tg-FRT cells to confirm the increase in Tg secretion. Cells were transfected with 25nM siRNAs for 36 h, media exchanged and conditioned for 8 h, Tg-FT was immunoprecipitated from media samples, and Tg-FT amounts were analyzed via immunoblotting. Multiple individual siRNAs recapitulated the increase in C1264R secretion.

**Figure EV8.**
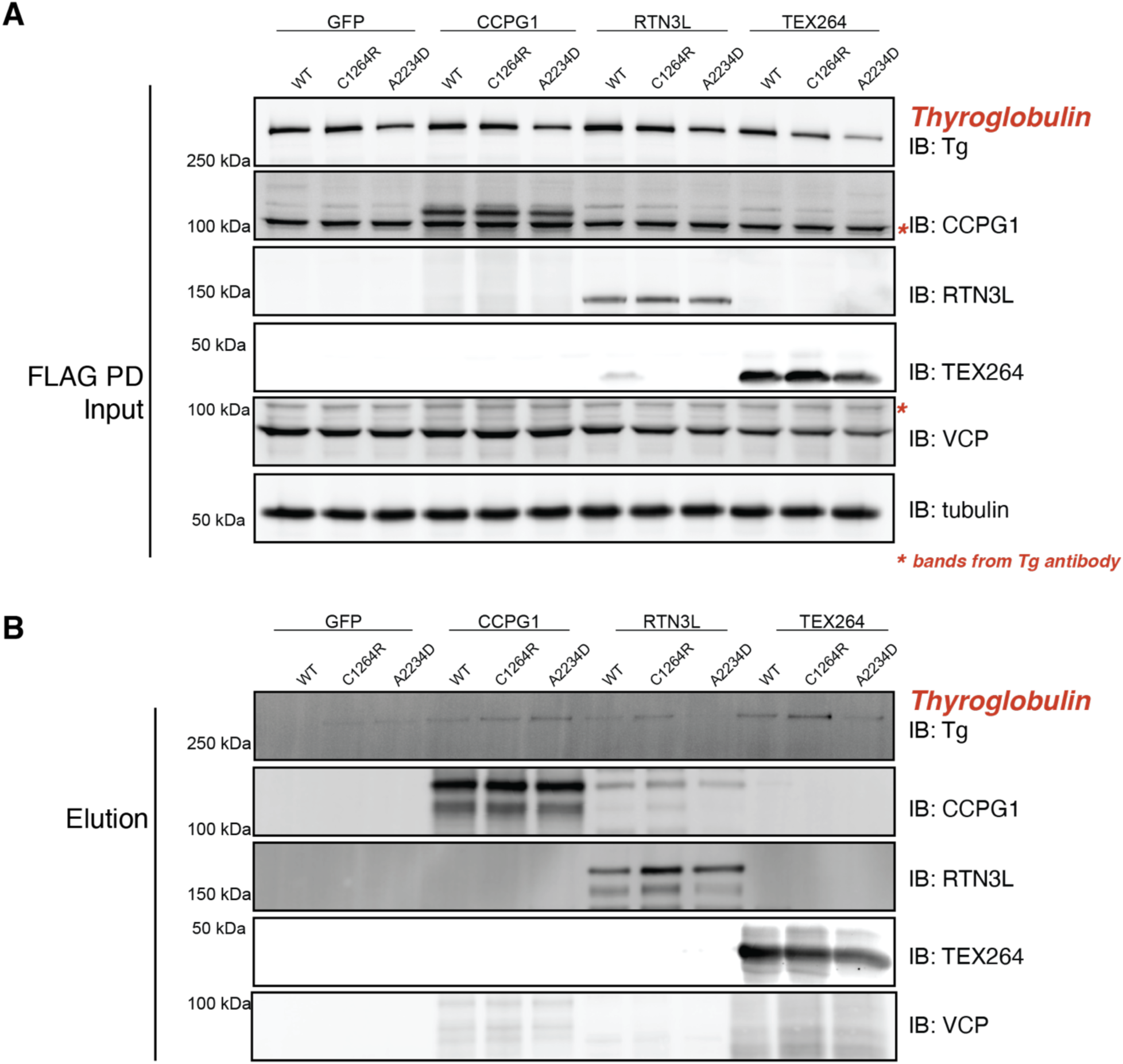
Mutant Tg Shows Selective Enrichment with TEX264 in a Screen of Multiple ER-phagy Receptors. (A) Western blot of FLAG-tagged ER-phagy receptors in cells expressing Tg. (B) FLAG co-IP elution samples. Western blot shows enrichment of ER-phagy receptor after pulldown (IB: CCPG1, RTN3L, TEX264). Selective enrichment of mutant Tg is present with TEX264 expression. The selective enrichment is not detectable in receptors CCPG1, RTN3L, or SEC62 (probed in Fig 6). Additionally, VCP shows no interaction with TEX264, suggesting that its interaction with Tg may be independent of ER-phagy receptors.

**Figure EV9.**
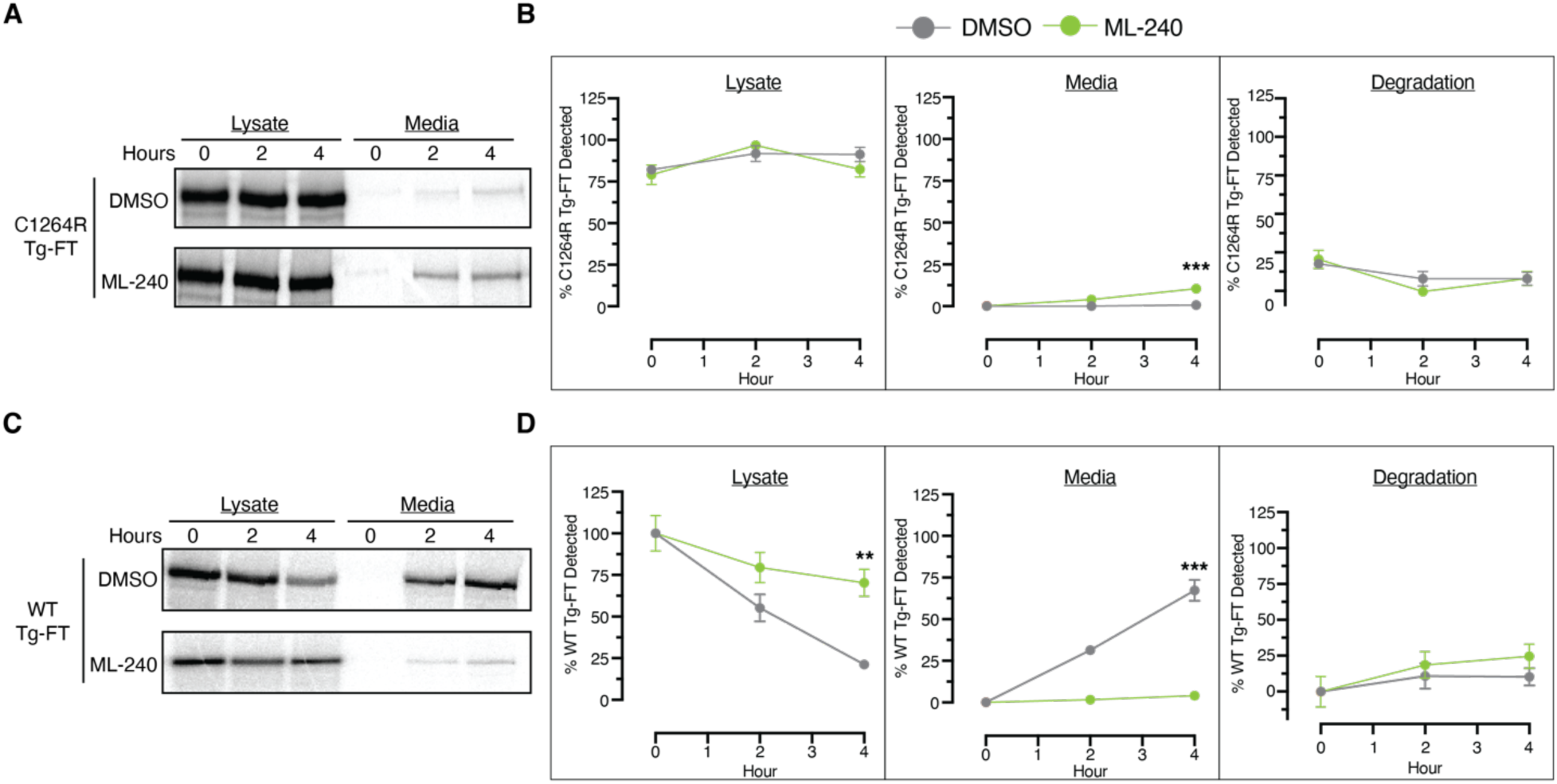
Pulse chase analysis of WT and C1264R Tg with pharmacological VCP inhibition. Autoradiographs and quantifications of pulse-chase analysis of C1264R Tg-FT (A-B) and WT Tg-FT (C-D) in FRT cells with ML-240 treatment. Cells were pre-treated with ML-240 or DMSO for 15 minutes prior to pulse labeling with EasyTag ^35^S Protein Labeling Mix (Perkin Elmer, NEG772007MC) for 30 minutes and chased for 4 hours with DMSO or ML-240 treatment, collecting samples at 0-, 2-, and 4-hour timepoints. Autoradiographs from a representative experiment are shown in A & C. Quantification is shown in C & D. Data is normalized to timepoint of maximum Tg recovery and represented as mean ± SEM. Statistical testing performed using an unpaired student’s t-test with Welch’s correction, *p<0.05, **p<0.005, ***p<0.0005.

**Figure EV10.**
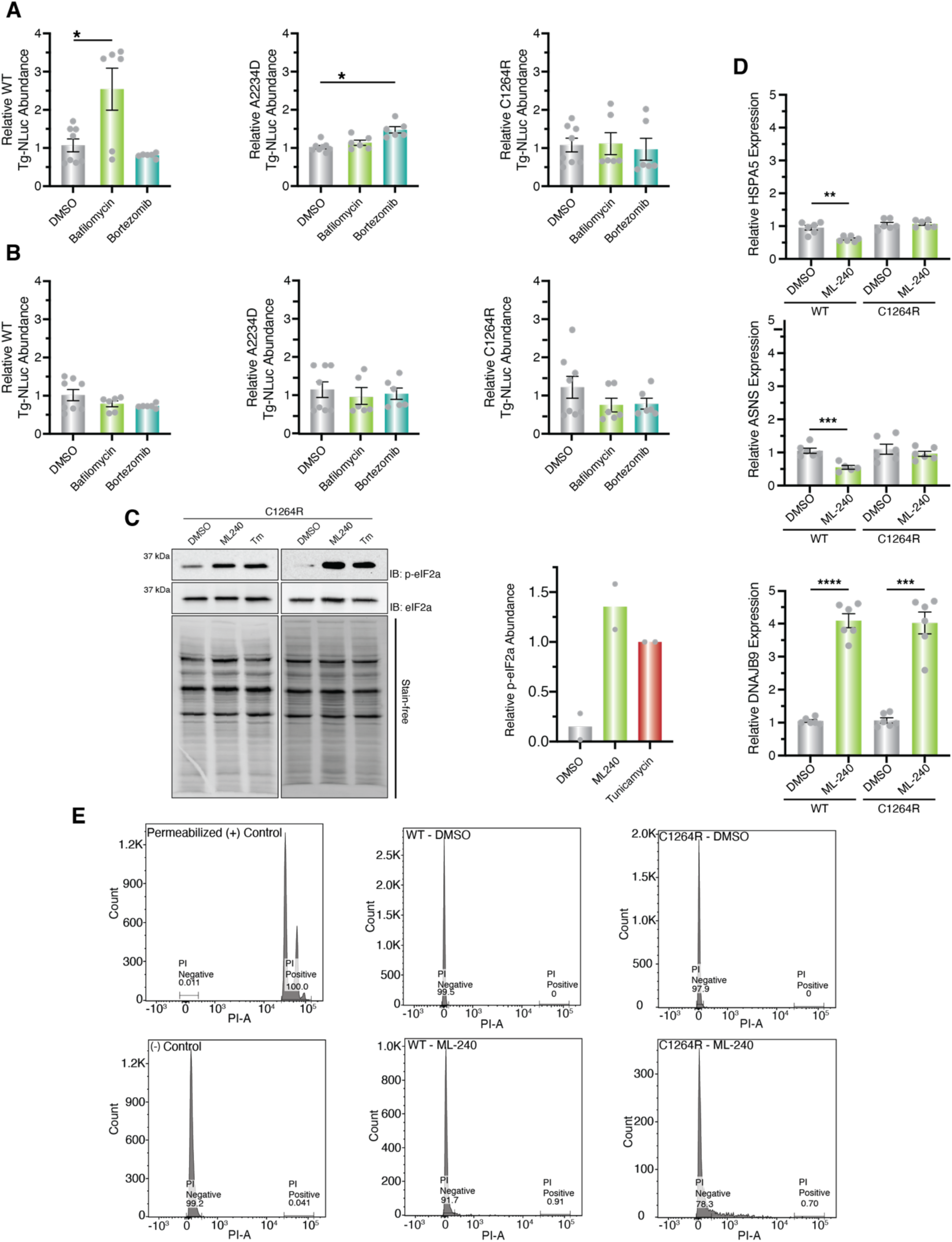
TRIP of C1264R Tg-FT FRT cells with pharmacological VCP inhibition. (A – B) Effect of pharmacologic inhibition of protein degradation in Tg retention and secretion. HEK293T cells stably expression Tg-NanoLuc variants were treated with proteasomal degradation inhibitor Bortezomib (10 µM) or lysosomal inhibitor Bafilomycin A1 (10 µM) for 8 hours. Prior to treatment, the media was exchanged to condition secreted Tg. Tg lysate amounts (A) or media amounts (B) were then measured by the nano-glo luciferase assay system. Data is normalized to DMSO condition and represented as mean ± SEM. Statistical testing performed using an unpaired student’s t-test with Welch’s correction, *p<0.05, **p<0.005, ***p<0.0005 (C – D) Assessment of UPR upregulation with ML-240 treatment. (C) Western blot analysis to assess phospho-eIF2α levels in of C1264R Tg-FT FRT cells treated with VCP inhibitor ML240 (10 µM), tunicamycin (1 µg/mL), or vehicle (0.1% DMSO) for 2 hours. Lysate samples were analyzed via immunoblotting. Data is normalized to the mean C1264R Tg-FT abundance of tunicamycin-treated samples. Data represented as mean ± SEM. (D) Activation of UPR markers monitored via qPCR in C1264R Tg-FT FRT cells treated with ML-240 (10μM) for 3 hours. HSPA5 and ASNS expression remained unchanged in C2164R Tg-FT FRT cells but led to a significant decrease in WT Tg-FT FRT cells. Only DNAJB9 showed a significant increase in transcript levels for both C1264R and WT Tg-FT FRT cells. This suggest that ML-240 dependent rescue of C1264R Tg is not due to global remodeling of the ER proteostasis network via UPR activation. Data was first normalized to a GAPDH loading control followed by normalization to median expression of DMSO treated samples and represented as mean ± SEM. Statistical testing performed using an unpaired student’s t-test with Welch’s correction, *p<0.05, **p<0.005, ***p<0.0005. Primers for detection described in **Table EV2**. (E) Viability analysis using Propidium iodide control samples, WT & C164R Tg-FT FRT cells with ML-240 treatment. Cells were treated with DMSO or ML-240 (10μM) for 4 hours, harvested, and stained with propidium iodide (1 μg/mL). FRT cells permeabilized with 0.2% Triton were used as a positive staining control. Unstained, non-permeabilized FRT cells were used as a negative staining control. Permeabilized and unstainined, non-permeabilized FRT cells described in were used to assess cell viability of ML-240 treated WT and C1264R Tg-FT FRT cells.

