## Appendix for "Time-Resolved Interactome Profiling Deconvolutes Secretory Protein Quality Control Dynamics"

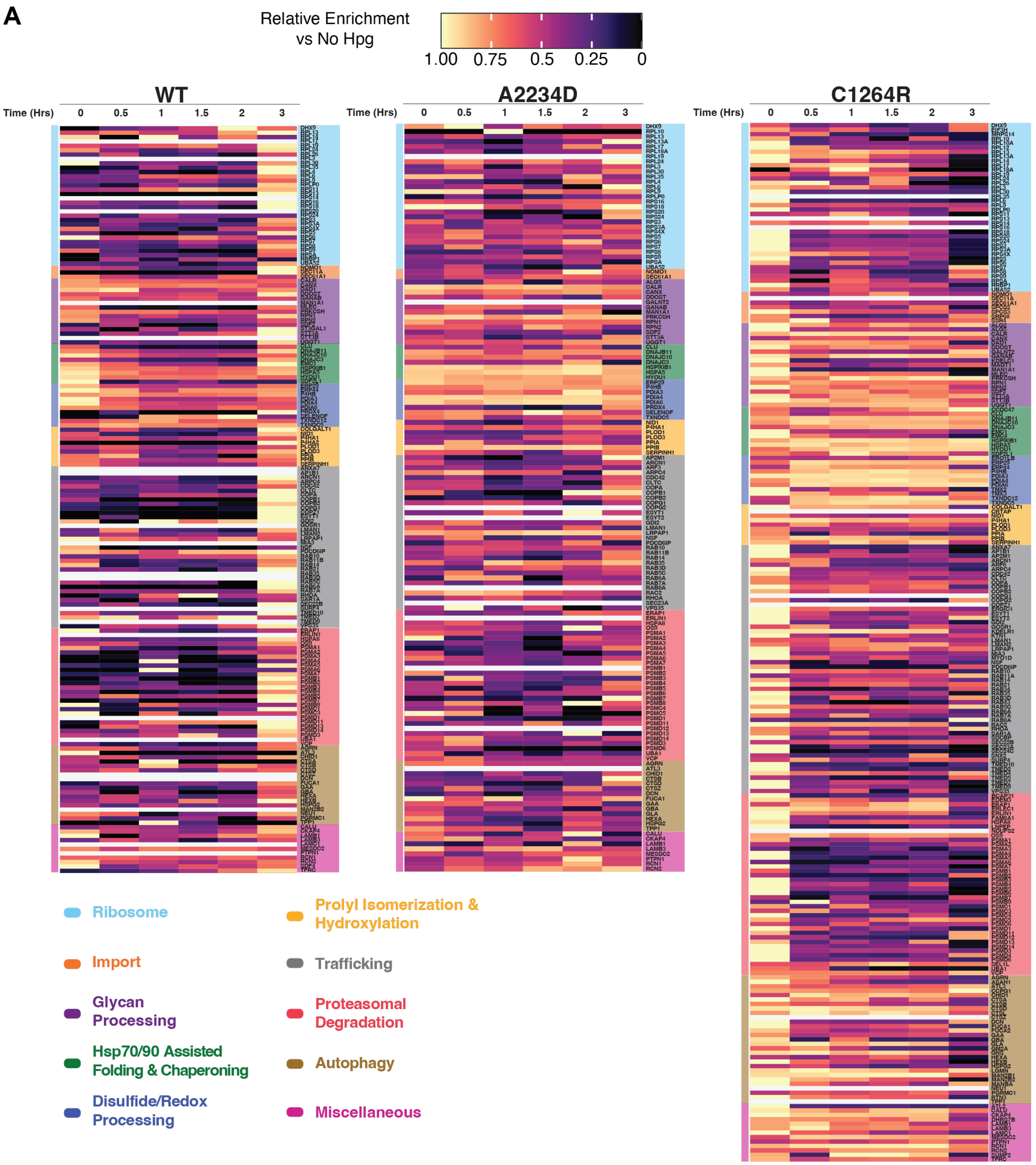
A
Relative Enrichment
vs No Hpg
1.00 0.75 0.5 0.25 0
WT
A2234D
C1264R
Time (Hrs)
0
0.5
1
1.5
2
3
Ribosome
Import
Glycan Processing
Hsp70/90 Assisted Folding & Chaperoning
Disulfide/Redox Processing
Prolyl Isomerization & Hydroxylation
Trafficking
Proteasomal Degradation
Autophagy
Miscellaneous

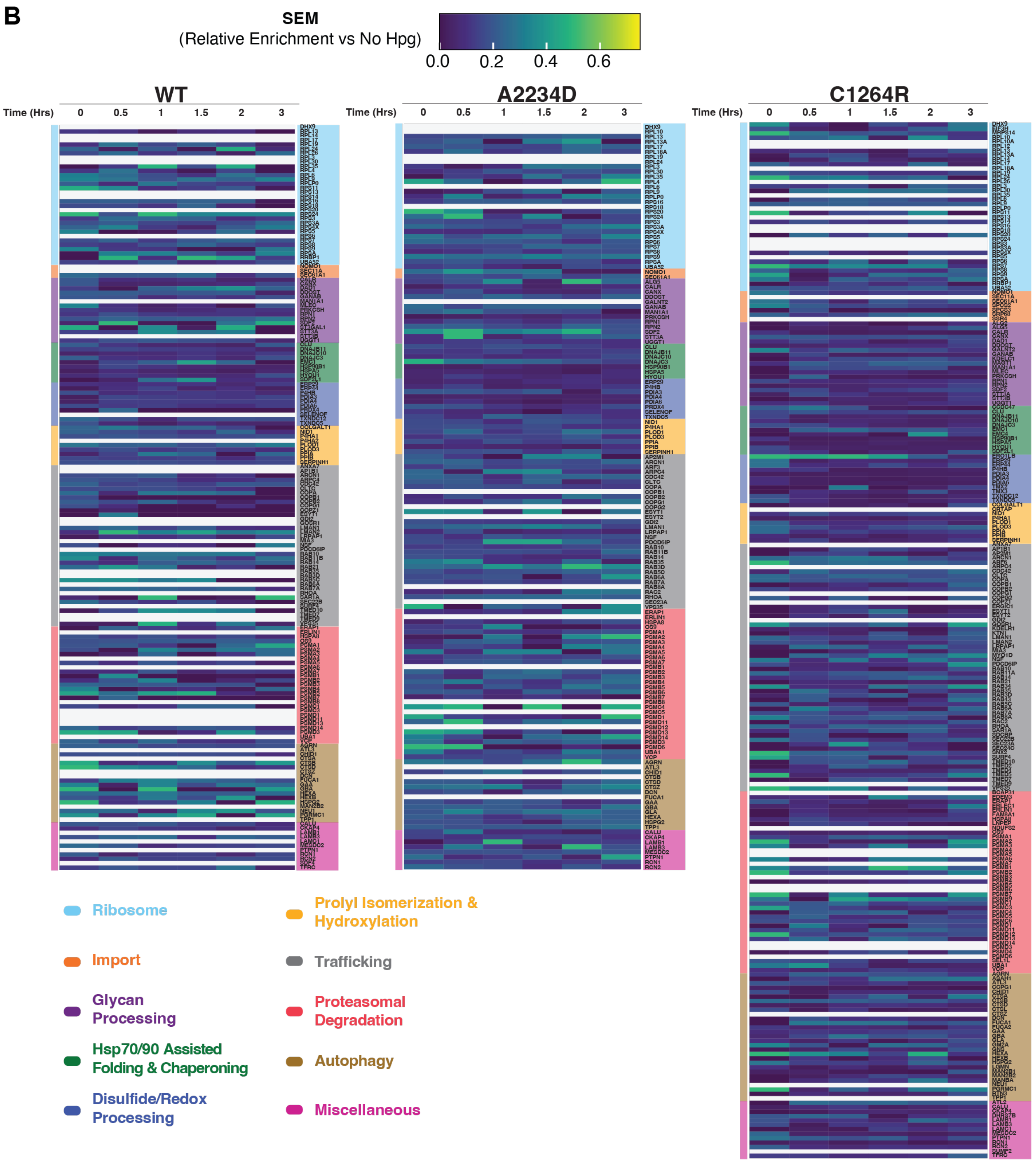
B
SEM
(Relative Enrichment vs No Hpg)
0.0
0.2
0.4
0.6
WT
A2234D
C1264R
Time (Hrs)
0
0.5
1
1.5
2
3
Ribosome
Import
Glycan Processing
Hsp70/90 Assisted Folding & Chaperoning
Disulfide/Redox Processing
Prolyl Isomerization & Hydroxylation
Trafficking
Proteasomal Degradation
Autophagy
Miscellaneous

**Appendix Figure S1 – Summary of Tg TRIP Data –scaled heatmap**

(A) Analysis showing the scaled log2 fold change enrichment of Tg interactors measured by TRIP for time-resolved analysis. Chase samples were pulse-labeled with Hpg (200µM) for 1 hour. Cells were harvested at specified time points and cross-linked with DSP (0.5mM) for 10 minutes to capture transient proteoastasis network interactions. Lysates were functionalized with TAMRA-Azide-PEG-Desthiobiotin probe CuAAC Click reaction. Chase samples were processed through the dual affinity purification TRIP workflow and processed for mass spectrometry. (-) Hpg samples were processed through the entire dual affinity purification TRIP workflow including 3-hour chase period, absent Hpg labeling, and used for enrichment analysis. Data were processed in R with custom scripts. TMT abundances across chase samples were normalized to Tg TMT abundance as described in the Materials and Methods section of the manuscript. For relative enrichment analysis, the means of log2 interaction differences were scaled to values from 0 to 1, where a value of 1 represented the time point at which the enrichment reached the maximum, while log2 values below the (-) Hpg condition were set to zero.

(B) Standard error of the mean (SEM) of the scaled log2 fold change enrichment of Tg interactors. n = 5 for WT and n= 6 for A2234D and C1264R.

Source data can be found in **Dataset EV4**. Script available at github.com/wrighmt1/2022_TRIP.

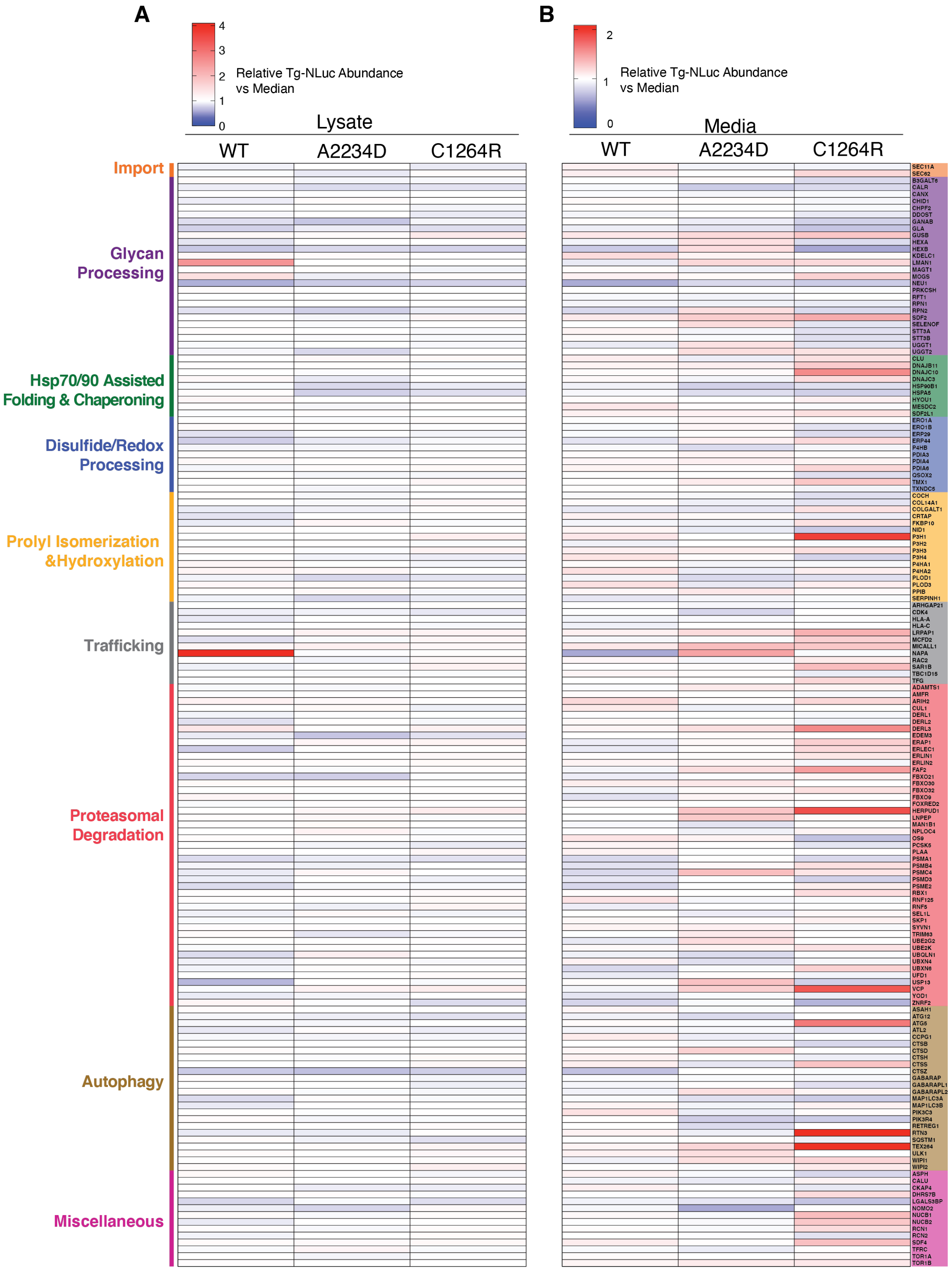

A
4
3
2
1
0
Relative Tg-NLuc Abundance vs Median
Lysate
WT
A2234D
C1264R
Import
Glycan Processing
Hsp70/90 Assisted Folding & Chaperoning
Disulfide/Redox Processing
Prolyl Isomerization &Hydroxylation
Trafficking
Proteasomal Degradation
Autophagy
Miscellaneous
B
2
1
0
Relative Tg-NLuc Abundance vs Median
Media
WT
A2234D
C1264R
SEC11A
SEC62
B3GALT6
CALR
CANX
CHID1
CHPF2
DDOST
GANAB
GLA
GUSB
HEXA
HEXB
KDELC1
LMAN1
MAGT1
MOGS
NEU1
PRKCSH
RFT1
RPN1
RPN2
SDF2
SELENOF
STT3A
STT3B
UGGT1
UGGT2
CLU
DNAJB11
DNAJC10
DNAJC3
HSP90B1
HSPA5
HYOU1
MESDC2
SDF2L1
ERO1A
ERO1B
ERP29
ERP44
P4HB
PDIA3
PDIA4
PDIA6
QSOX2
TMX1
TXNDC5
COCH
COL14A1
COLGALT1
CRTAP
FKBP10
NID1
P3H1
P3H2
P3H3
P3H4
P4HA1
P4HA2
PLOD1
PLOD3
PPIB
SERPINH1
ARHGAP21
CDK4
HLA-A
HLA-C
LRPAP1
MCFD2
MICALL1
NAPA
RAC2
SAR1B
TBC1D15
TFG
ADAMTS1
AMFR
ARIH2
CUL1
DERL1
DERL2
DERL3
EDEM3
ERAP1
ERLEC1
ERLIN1
ERLIN2
FAF2
FBXO21
FBXO30
FBXO32
FBXO9
FOXRED2
HERPUD1
LNPEP
MAN1B1
NPLOC4
OS9
PCSK5
PLAA
PSMA1
PSMB4
PSMC4
PSMD3
PSME2
RBX1
RNF125
RNF5
SEL1L
SKP1
SYVN1
TRIM63
UBE2G2
UBE2K
UBQLN1
UBXN4
UBXN6
UFD1
USP13
VCP
YOD1
ZNRF2
ASAH1
ATG12
ATG5
ATL2
CCPG1
CTSB
CTSD
CTSH
CTSS
CTSZ
GABARAP
GABARAPL1
GABARAPL2
MAP1LC3A
MAP1LC3B
PIK3C3
PIK3R4
RETREG1
RTN3
SQSTM1
TEX264
ULK1
WIPI1
WIPI2
ASPH
CALU
CKAP4
DHRS7B
LGALS3BP
NOMO2
NUCB1
NUCB2
RCN1
RCN2
SDF4
TFRC
TOR1A
TOR1B

**Appendix Figure S2 - Summary of siRNA screening data**

(A) Analysis showing the relative Tg-NLuc abundance changes in lysate with siRNA knockdown of select genes. Approximately 36 hours after transfection with 25nM siRNAs cells were replenished with fresh media and Tg-NLuc abundance in lysate was measure after 4 hours using the nano-glo luciferase assay system. Data was median normalized across individual 96-well plates (Chung et al., 2008). Data represents two independent experiments for WT-NLuc and A2234D-NLuc, and three independent experiments for C1264R NLuc. Cutoff criteria for hits were set to those genes that increased or decreased Tg-NLuc abundance in lysate or media by 3σ.

(B) Analysis showing the relative Tg-NLuc abundance changes in media with siRNA knockdown of select genes. Approximately 36 hours after transfection with 25nM siRNAs cells were replenished with fresh media and Tg-NLuc abundance in media was measure after 4 hours using the nano-glo luciferase assay system. Data was median normalized across individual 96-well plates (Chung et al., 2008). Data represents two independent experiments for WT-NLuc and A2234D-NLuc, and three independent experiments for C1264R NLuc. Cutoff criteria for hits were set to those genes that increased or decreased Tg-NLuc abundance in lysate or media by 3σ. Source data can be found in **Dataset EV5**.

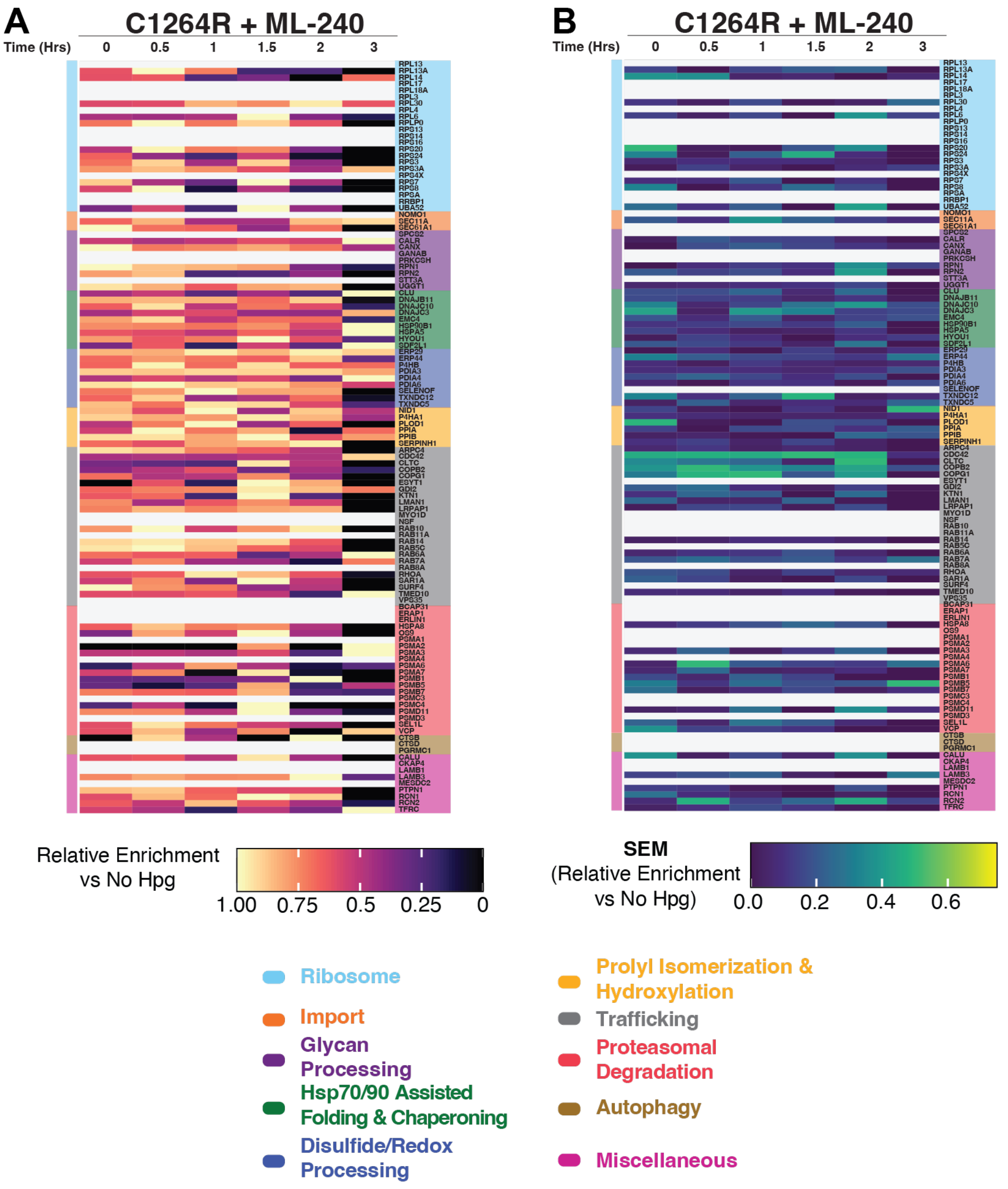


**Appendix Figure S3 -TRIP of C1264R Tg-FT FRT cells with pharmacological VCP inhibition**

(A) Analysis showing the scaled log2 fold change enrichment of C1264R Tg interactors measured by TRIP with ML-240 treatment. Chase samples were pulse labeled with Hpg (200 µM final concentration) in the presence of ML-240 (10 µM) for 1 h. Cells were dosed with ML-240 (10µM) throughout the chase period, harvested at specified time points and cross linked with DSP (0.5mM) for 10 minutes to capture transient proteoastasis network interactions. Lysates were functionalized with TAMRA-Azide-PEG-Desthiobiotin probe using copper catalyzed azide-alkyne cycloaddition (CuAAC). Chase samples were processed through the dual affinity purification TRIP workflow and processed for mass spectrometry. (-) Hpg samples were processed through the entire dual affinity purification TRIP workflow in the presence of ML-240 (10µM) including 3 h chase period, absent Hpg labeling, and used for enrichment analysis.

Data were processed in R with custom scripts. TMT abundances across chase samples were normalized to Tg TMT abundance as described within the Materials and Methods section of the manuscript. For relative enrichment analysis, the means of log2 interaction differences were scaled to values from 0 to 1, where a value of 1 represented the time point at which the enrichment reached the maximum, while log2 values below the (-) Hpg condition were set to zero.

(B) Analysis showing the SEM of the scaled log2 fold change enrichment of C1264R Tg interactors measured by TRIP with ML-240 treatment. Chase and (-) Hpg samples were processed as described above in (A). Data were processed in R with custom scripts. TMT abundances across chase samples were normalized to Tg TMT abundance as described within the Materials and Methods section of the manuscript. Standard error of the mean (SEM) was then calculated from these enrichment values to examine the reproducibility of these measurements. Script available at github.com/wrighmt1/2022_TRIP. Source data for (A) – (B) can be found in **Dataset EV4.**
